# Neuronal resonance can be generated independently at distinct levels of organization

**DOI:** 10.1101/2020.05.26.117309

**Authors:** Eran Stark, Horacio G. Rotstein

## Abstract

Neuronal resonance is defined as maximal amplification of the response of a system to a periodic input at a finite non-zero input frequency band. Resonance has been observed experimentally in the nervous system at the level of membrane potentials, spike times, post-synaptic potentials, and neuronal networks. It is often assumed that resonance at one level of organization endows resonance at another level, but how the various forms of neuronal resonances interact is unknown. Here we show that a direct link of the frequency response properties across neuronal levels of organization is not necessary. Using detailed biophysical modeling combined with numerical simulations, extracellular recordings, and optogenetic manipulations from behaving mice, we show how low-pass filtering, high-pass filtering, and amplification mechanisms can generate resonance at a single level of organization. Subthreshold resonance, synaptic resonance, and spiking resonance can each occur in the lack of resonance at any other level of organization. In contrast, frequencydependent mechanisms at several levels of organization are required to generate the more complex phenomenon of network resonance. Together, these results show that multiple independent mechanisms can generate resonance in neuronal systems.

## INTRODUCTION

Resonance refers to the maximal response of a system to oscillatory inputs at a finite non-zero (“resonant”) frequency. The concept originated in electric (capacitor-inductor) and mechanical (spring-mass) linear systems subjected to an external forcing input. In the absence of resistance or friction, the response of such a system to oscillatory inputs is maximal when the input frequency matches the natural frequency of the unforced oscillator; in such a case, the resonant and natural frequencies are identical. In the presence of resistance, when the unforced system exhibits damped oscillations, the response variable exhibits a peak amplitude at a resonant frequency that is close to but distinct from the natural frequency. While resonance is often associated with (sustained or damped) oscillators, it is a property of more general systems. In fact, linear 2D systems that do not display intrinsic oscillations (i.e. do have a stable node) can exhibit resonance for a wide range of parameter values (Richardson et al., 2003; Rotstein and Nadim, 2014). Typically, these systems exhibit an overshoot in response to step function inputs.

Because neuronal systems are electric circuits, it is expected that they will exhibit resonance. Indeed, neuronal resonance has been observed at multiple levels of organization and quantified using various metrics, in all cases capturing the notion of optimal gain. In the simplest case, similarly to RLC circuits, the subthreshold (membrane potential) response of an isolated neuron to oscillatory inputs has been measured in terms of the impedance *Z* amplitude profile (ZAP), quantifying the amplitude response as a function of the input frequency *f* (Gutfreund et al., 1995; Hutcheon et al., 1996; Hu et al., 2002, 2009; Hutcheon and Yarom, 2000; Puil et al., 1986; Wang, 2010). The impedance is a complex quantity having amplitude and phase. A neuron exhibits subthreshold resonance if *|Z(f)|* peaks at a non-zero frequency. Otherwise, individual neurons may behave as low-pass filters (Puil et al., 1986; Pike et al., 2000; Zemankovics et al., 2010) or may exhibit more complex behavior depending on the number of ionic currents and their time scales (Richardson et al., 2003; Pike et al., 2000; Fox, Rotstein, and Nadim, 2016, SFN Abstract; Rotstein, 2017b). In addition to subthreshold resonance, spikes may occur at specific frequencies of an oscillatory input current (Pike et al., 2000). Moreover, at the level of synaptic transmission, the amplitude of post-synaptic potentials (PSPs) may peak at some instantaneous frequency of the pre-synaptic spikes (Markram et al., 1998; Izhikevich et al., 2003; Drover et al., 2007). Finally, resonance may occur at the network level (Stark et al., 2013): resonance has been observed in the spike timing of one group of neurons when an oscillatory depolarizing input was applied to another group of neurons.

Theoretical studies have shown that subthreshold resonance can be communicated to the spiking regime in the firing rate (Brunel and Hakim, 2003) and other measures of spiking activity (Engel et al., 2008; Rotstein, 2017c). A possible implication of this observation is that resonances communicate across levels of neuronal organization, either directly or indirectly. For instance, subthreshold resonance at theta frequencies may be expected to create spiking resonance at theta frequencies, which may in turn generate network resonance at theta frequencies when resonant spiking neurons interact with other neurons. Alternatively, the interplay of the positive and slower negative feedback effects across interacting levels of organization may communicate resonance across these levels. However, direct in vivo activation of hippocampal CA1 pyramidal cells that have been shown to exhibit subthreshold resonance in vitro did not produce network resonance (Stark et al., 2013). Moreover, modelling suggests that spiking resonance in subthreshold resonant neurons is restricted to relatively small input amplitudes (Engel et al., 2008; Rotstein, 2017c). Thus, it is still unclear whether and under what conditions resonance at one level of organization implies resonance at another level.

The specific question we address in this paper is whether resonance observed at one level of organization is necessarily an outcome of (i.e. inherited from) resonance at another level. An alternative is the existence of independent processes that may share some building blocks, and act to generate resonance at distinct levels. This alternative scenario does not preclude the existence of neuronal systems in which resonances are communicated across levels of organization. To tackle this question, we combine detailed biophysical modeling, numerical simulations, extracellular recordings, and optogenetic manipulations in freely-moving mice. We identify and analyze a number of case studies at various levels of organization (subthreshold, spiking, synaptic, network) where the generation of resonance depends on mechanisms intrinsic to each level.

## RESULTS

### Neuronal circuitry that exhibits intrinsic oscillations does not necessarily exhibit resonance at the same frequency

To define resonance in a way that encompasses the various levels of neuronal organization, consider a system for which the input is current or a current-related quantity (e.g. photocurrent), and the output is voltage or a voltage-related quantity (e.g. firing rates, LFP; **Fig. 1A**). In the context of rhythmic systems, one can differentiate between two types of responses: an oscillator and a resonator. In an electric oscillator (**Fig. 1A, left**), the input is a square pulse of current (no time-dependent dynamics), and the output is an oscillatory voltage. The second type of rhythmic system is a resonator (**Fig. 1A, right**). In its simplest form, a resonator responds to an oscillatory input current with an oscillatory output voltage that displays frequency-dependent amplitude peaking at a non-zero frequency.

**Figure 1.**
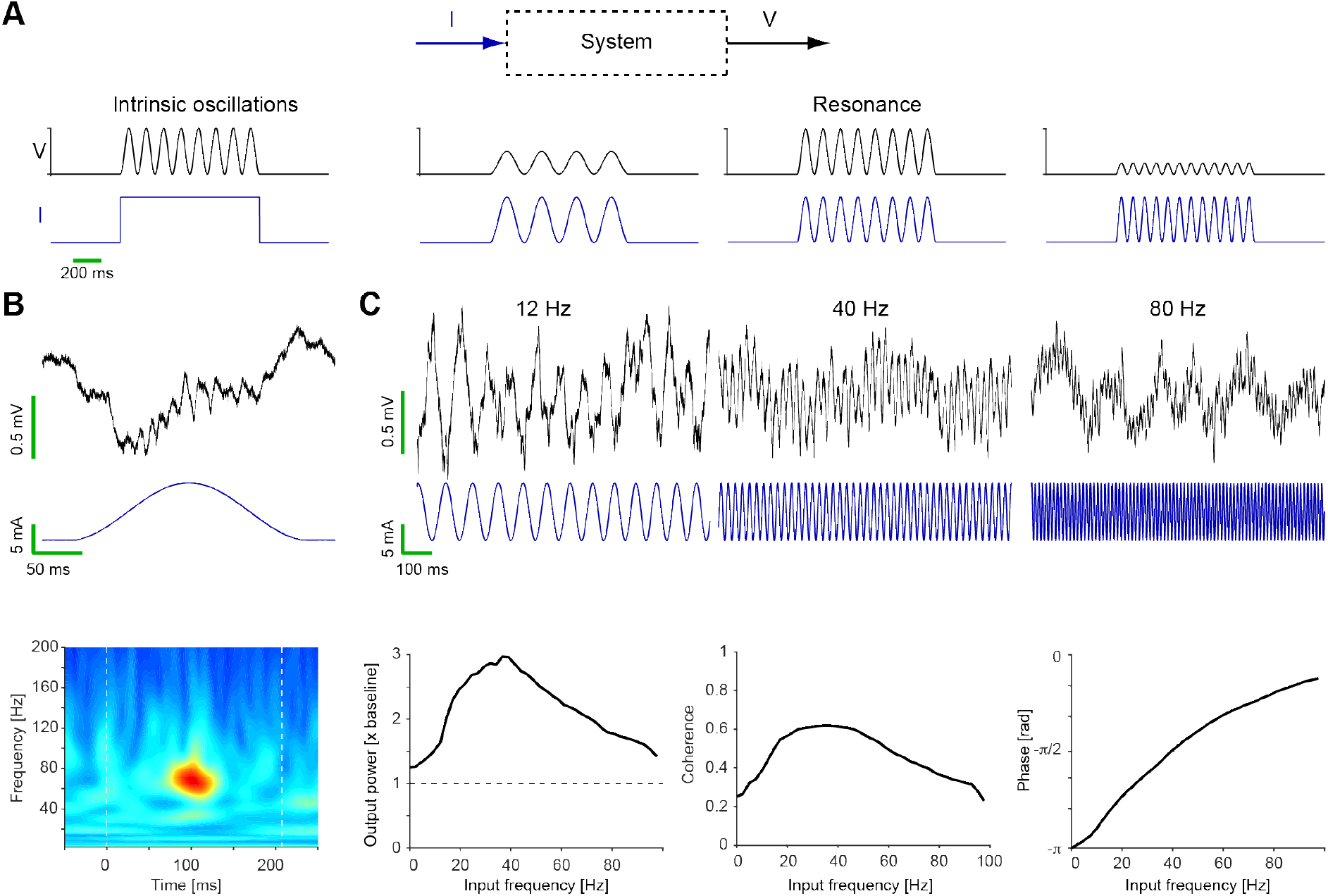
Resonance and intrinsic oscillations are distinct phenomena. (**A**) For a given system (top), *intrinsic oscillations* are defined as a single frequency rhythm following a non-periodic (e.g., DC) input (left). Thus, they are a property of the system and may be uncovered by the input. *Resonance* is defined as an oscillatory output with maximal amplitude at a single nonzero finite frequency band following an oscillatory input at multiple frequency bands (right). Resonators need not be intrinsic oscillators (persistent or damped). (**B**) Optically-induced intrinsic oscillations in CA1. Recordings are from a freely-moving CaMKII::ChR2 mouse. Wide-band traces (0.1-7,500 Hz) during illumination with a sinusoidal waveform (peak applied current 10 mA, corresponding to 0.01 mW/mm^2^ at the recording site). Focal illumination induces high-frequency oscillations in the local field potential (LFP), with a peak frequency of 65-70 Hz. Time-frequency decomposition shows average of 40 individual stimuli. (**C**) Optically-induced resonance in CA1. *Top:* wide-band traces show LFP (same recording site as in **B**) during segments of a chirp input. At low input frequencies, peak illumination induces maximal depolarization, corresponding to LFP negativity. *Bottom:* Output spectrum, input-output coherence, and input-output phase lag as a function of input frequency. The output power and coherence peak at 40 Hz. Thus, intrinsic oscillations and resonance in CA1 have distinct frequencies (65-70 vs. 40 Hz).

Mathematically, oscillations cannot be intrinsically generated in a one-dimensional system. In a 2D dynamical system, sustained oscillations correspond to a limit cycle in the phase-plane diagram, whereas damped oscillations correspond to a stable equilibrium (focus; Izhikevich, 2007). Resonance can be obtained in a 2D system even in the lack of intrinsic (i.e. sustained or damped) oscillations (Hutcheon et al., 1996; Brunel and Hakim, 2003; Rotstein, 2014; Rotstein and Nadim, 2014). There is a region in the parameter space in which both resonance and damped oscillations may coexist (Richardson et al., 2003). However, the resonant and intrinsic (natural) frequencies do not coincide (Richardson et al., 2003; Rotstein and Nadim, 2014). It has been argued that resonators and oscillators are different expressions of the same underlying mechanism (Lampl and Yarom, 1997) and that they belong to an oscillatory hierarchy where non-oscillatory resonators are at the base and oscillations emerge for high enough amplification levels (Hutcheon and Yarom, 2000). Indeed, in a deterministic 2D system, increasing levels of amplification may cause a transition from non-oscillatory resonators, to damped oscillators, to sustained oscillators (Hutcheon and Yarom, 2000). However, for 2D linear systems, damped oscillations may exist in the lack of resonance (Brunel and Hakim, 2003, Rotstein and Nadim, 2014). These results indicate that resonance and oscillations are related, but conceptually different.

Although permitted mathematically, to our knowledge there has been no prior experimental evidence of non-resonating neuronal oscillations. Thus, it is unclear whether an intact neuronal network that can generate intrinsic oscillations necessarily resonates at the same frequency. To test this, we implanted multi-site diode-probes in freely-moving CaMKII::ChR2 mice and activated pyramidal cells in CA1. Illumination had the form of a single cycle of a 5 Hz sinusoid, (i.e. 200 ms), with an intensity of 0.01 mW/mm^2^ at the recording site (0.5 μW at the fiber tip). Under these conditions, the local field potential (LFP) exhibited a slow wave of depolarization accompanied by high frequency oscillations (60-80 Hz; **Fig. 1B**; see also Stark et al., 2014, 2015). We then applied an oscillatory sweep of frequencies (linear chirp, from 0-100 Hz or from 100-0 Hz) at an identical intensity. At low input frequencies, the phase of the induced LFP response was opposite from the light, consistent with an extracellular signature of intra-cellular depolarization (**Fig. 1B** and **1C**). The amplitude of the induced LFP response initially increased and then decreased monotonically with increasing frequency, exhibiting a resonant peak around 40 Hz (**Fig. 1C**). This shows that LFP oscillations that emerge at the high-gamma (60-80 Hz) band are not accompanied by high-gamma LFP resonance. Thus, an intrinsic neuronal oscillator is not necessarily a resonator.

### A passive membrane with a single resonant current yields subthreshold resonance

From this point on, we focus exclusively on neuronal resonators. Since resonance has been observed in the membrane potential of individual neurons (Gutfreund et al., 1995; Hutcheon et al., 1996; Hutcheon and Yarom, 2000), we begin by revisiting resonance in simple neuronal models. A passive membrane can be modelled as a parallel RC circuit with a battery in series to the resistor (**Fig. 2A**). When oscillatory input current (*I*) of various frequencies (*f*) is applied to such a circuit, the voltage output (*V*) behaves as a low-pass filter: the complex impedance, defined as *Z(f)=V(f)/I(f)*, exhibits an amplitude profile that decreased monotonically with frequency. The phase of the output relative to the input is also frequency dependent, and increases monotonically with the frequency, indicating a delayed response. This is exactly analogous to the Bode diagram of a parallel RC circuit, acting as a low-pass filter (LPF).

**Figure 2.**
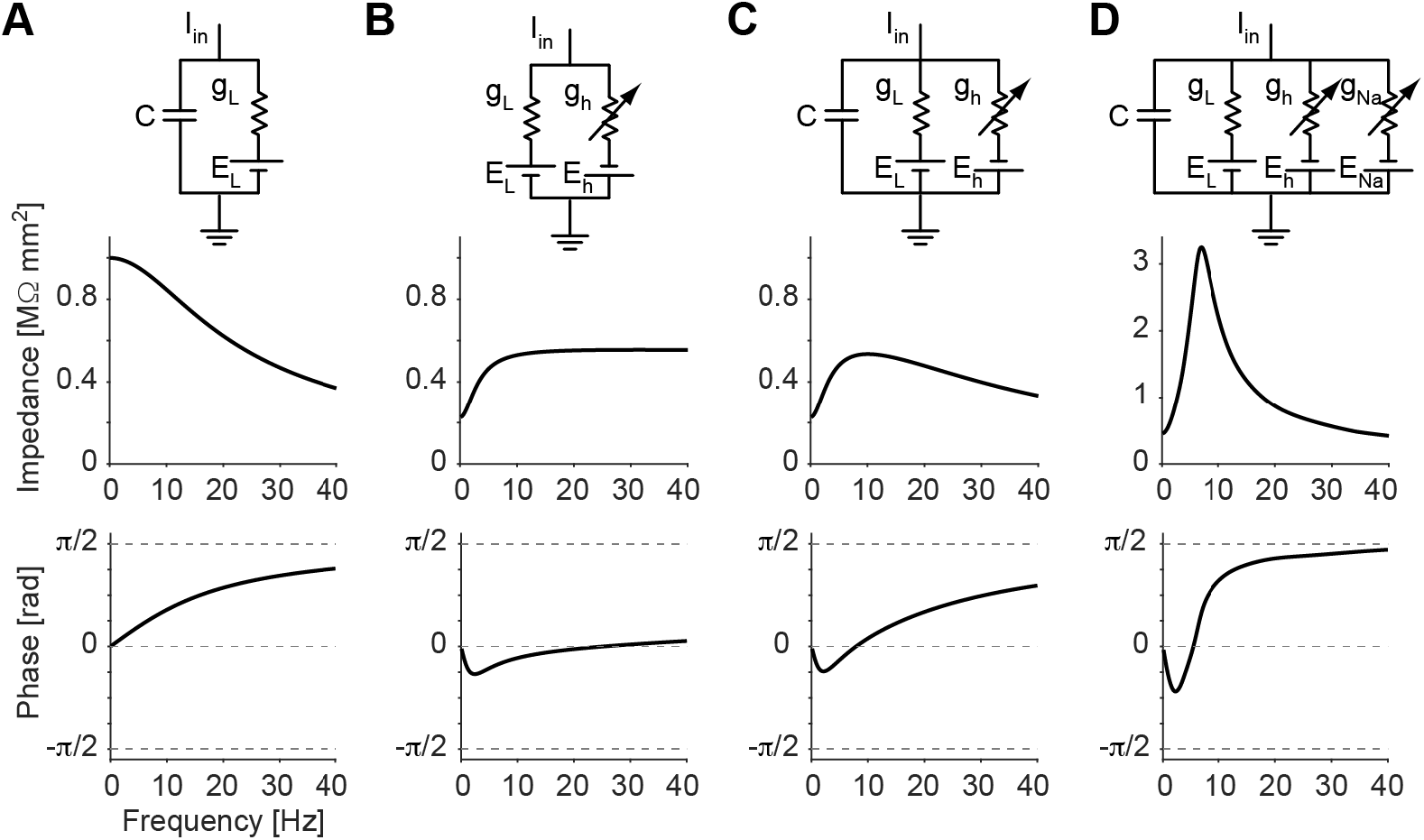
Subthreshold resonance can be generated by resonant currents acting as high-pass filters. (**A**) A leaky membrane (resistive and capacitive, i.e. an RC circuit) functions as a low-pass filter (LPF) of oscillatory input currents. (**B**) A resistive “membrane” with an inductive (“resonant”) current I_h_ acts as an RL network, displaying a high-pass filter (HPF) response to oscillatory inputs. (**C**) A leaky membrane with I_h_ acts as an RLC circuit and functions as a band-pass filter (BPF). (**D**) Adding an amplifying current (I_Na,p_) to a passive membrane with I_h_ yields a BPF with higher and sharper peak, resulting in a higher Q-factor (defined as the resonant frequency peak divided by the full-width at half-maximum).

In electrical circuits, a high-pass filter (HPF) can be obtained by replacing the capacitor with an inductor (a coil), yielding a parallel RL circuit. In the context of neuronal systems, a similar effect is obtained by minimizing capacitance and adding a current that opposes voltage changes; in this case, we used a hyperpolarization-activated sodium current (I_h_; **Fig. 2B**). Indeed, when the circuit receives an array of input frequencies, it exhibits a high-pass filter behavior with a phase advance in a subset of frequencies (negative phase). In addition to DC, the phase is exactly zero (i.e. the input current and output voltage are exactly synchronized in phase) at some non-zero frequency, sometimes referred to as the “phasonant” frequency (Rotstein, 2017b).

When I_h_ (a high-pass filter) is added to the passive membrane (a low-pass filter), we obtain the equivalent of a resistor-inductor-capacitor (RLC) circuit. This circuit behaves as the time-domain convolution of the individual RL and RC filters (multiplication in the frequency domain) yielding a bandpass filter (BPF; **Fig. 2C**). The dynamics remain quasi-linear around the equilibrium (Rotstein, 2017a). When voltage changes are further amplified, implemented here by adding a persistent sodium current (I_Na,p_), the hump in the frequency response becomes sharper (**Fig. 2D**) as stronger nonlinearities develop (Rotstein, 2017a; Rotstein 2015). Qualitatively, the outputs of the two resonant systems (the passive membrane with I_h_; and the passive membrane with I_h_ and I_Na,p_) in these regimes are similar. However, since the persistent sodium amplification is voltage-dependent and favors changes in voltage, I_Na,p_ contribution is not flat in the frequency domain and accentuates the resonant peak via positive feedback.

In sum, a 2D neuronal model of a passive membrane with a resonant current may exhibit resonance (**Fig. 2C-D**). This is equivalent to a passive RLC circuit that exhibits band-pass filtering properties. In general, the cutoff frequency of the HPF must be lower than the cutoff frequency of the LPF; otherwise, the filter will reject all frequencies (band-stop filter). Likewise, the neuronal subthreshold HPF, I_h_, must be slow enough compared to the membrane time constant in order for subthreshold resonance to emerge. Estimates of membrane time constants are on the order of 10 ms (R=1 MΩ mm^2^; C=10 nF/mm^2^; Dayan and Abbott, 2001), and estimates of τ_h_’s are around 45-90 ms (Poolos et al., 2002), indeed corresponding to a lower cutoff frequency.

### Spiking resonance is observed in intact animals and can occur without subthreshold resonance

Resonance can also be observed at the level of spiking. This has been shown in individual cortical neurons in vitro at the same frequency that induced subthreshold oscillations, when the amplitude of the input current was slightly increased (Hutcheon et al., 1996). Spiking resonance has also been observed in hippocampal slices, where the extracellular spiking of fast-spiking interneurons was confined to beta-gamma (10-50 Hz) input frequencies (Pike et al., 2000). To determine whether spiking resonance can also be obtained in other preparations, we applied linear chirps to parvalbumin-immunoreactive cells in PV::ChR2 mice implanted with multi-site diode probes. Some cells exhibited a resonant peak in the gamma range (**Fig. 3A**), demonstrating that spiking resonance can be observed in intact animals.

**Figure 3.**
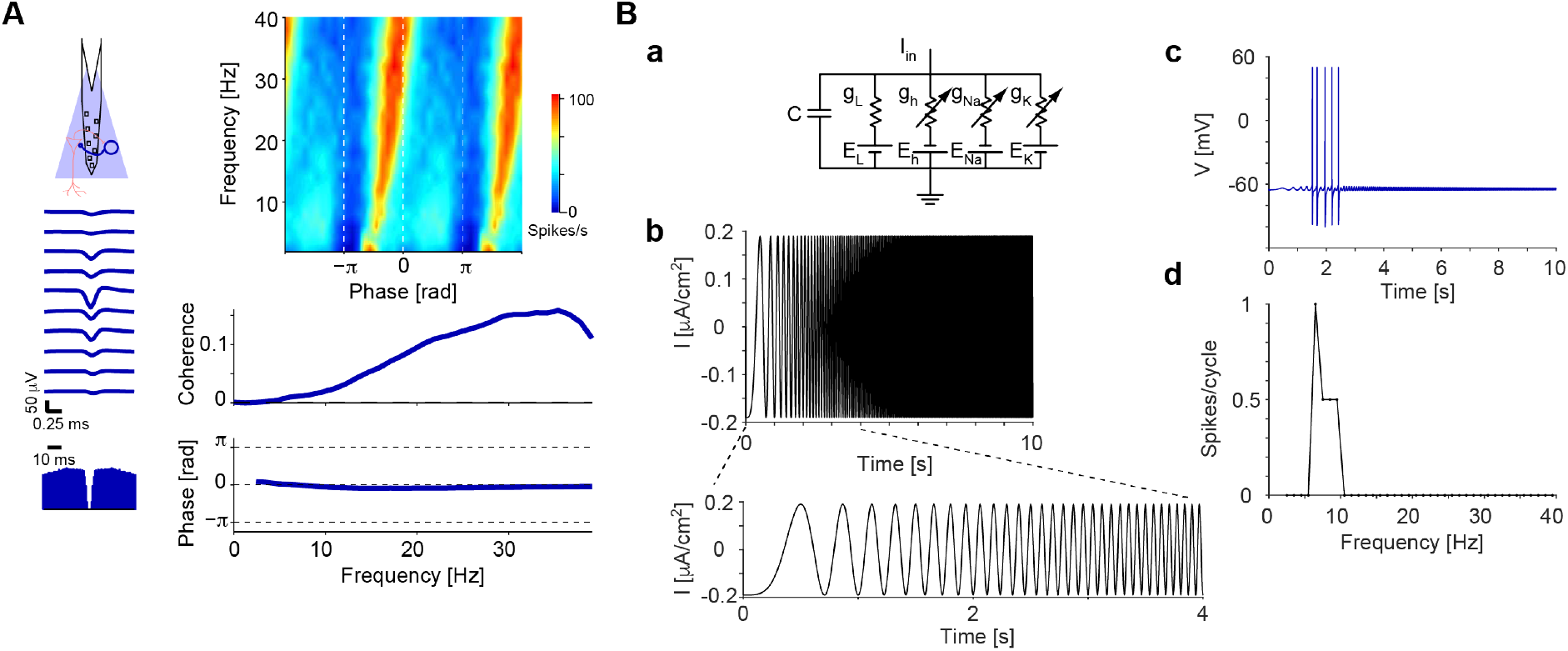
Single neurons can exhibit spiking resonance. (**A**) Gamma-band spiking resonance in a biological neuron. A parvalbumin-immunoreactive (PV) cell in CA1 of a PV::ChR2 mouse was illuminated with a linearly time-varying frequency (chirp) pattern. The cell spiked at all input frequencies, exhibiting a peak at the gamma range (30-40 Hz). (**B**) Theta-band spiking resonance in a model neuron. In the presence of subthreshold resonance, a single neuron can exhibit spiking resonance for low yet supra-threshold input amplitudes. (**a**) Circuit diagram of the conductance-based model of the Hodgkin-Huxley type with spiking currents. (**b**) The injected current pattern. (**c**) Membrane potential follows the injected current, crossing threshold at a non-zero frequency. (**d**) Quantification of spiking resonance by the number of spiking per cycle (a point-process equivalent of the impedance).

One way to model spiking resonance is to add spiking currents to a model that exhibits subthreshold resonance. A minimal model in this case is four-dimensional, involving dynamics on the membrane potential and fast (transient) sodium, delayed-rectifier potassium, and h-currents (**Fig. 3Ba**). When such a model was perturbed with an oscillatory input current (**Fig. 3Bb**), spikes occurred only in a narrow range of frequencies (**Fig. 3Bc**). This band-limited spiking can be quantified by the number of spikes observed at each input frequency cycle (**Fig. 3Bd**). Thus, in such a conductance-based model of a spiking neuron, spiking resonance is completely inherited from the subthreshold resonant mechanisms. This is consistent with the firing rate resonance reported previously (Brunel and Hakim, 2003; Rotstein, 2017c). However, subthreshold and spiking regime resonance are not necessarily generated by identical mechanisms. To demonstrate this, it is sufficient to show that subthreshold resonance can be accompanied by spikes in cells that do not exhibit spiking resonance. To demonstrate a double dissociation, it is also required to show that spiking resonance can occur in the lack of subthreshold resonance.

We began with the first direction, by asking whether subthreshold resonance may be accompanied by spiking activity, but without spiking resonance. We used the same 4D model and applied oscillatory current inputs identical in their frequency profile (0-40 Hz linear chirps; **Fig. 3Bb**) but at various amplitudes. In all tested cases, the membrane potential yielded a peak at a non-zero finite frequency (**Fig. 4A**), indicating subthreshold resonance. The resonant frequency was similar across all tested input amplitudes (8-10 Hz; green line in **Fig. 4A**). At low input amplitudes, no spikes were induced (**Fig. 4B**). At intermediate input amplitudes, spikes occurred at the peaks of the current input cycles, around the subthreshold resonant frequency (green lines in **Fig. 4B**). At the highest amplitudes, the largest number of spikes per cycle occurred at the lowest input frequencies tested (**Fig. 4B, right**). Thus, while at low input currents resonance affects spiking almost directly, at higher input currents the dominant property that is transmitted to the supra-threshold domain is the low-pass filter (LPF; **Fig. 2A**), stemming mainly from the passive properties of the membrane (RC) and shifting the resonant frequency towards DC. This corresponds to the fact that increasing the input amplitude causes an upward shift of the subthreshold impedance profile. For large enough input amplitudes, the impedance at DC (the resistance) will be above the spiking threshold. The asymmetric form of the subthreshold impedance profile (**Fig. 2C-D**) predicts a transition from resonance to a low-pass spiking response, as opposed to a transition to a wide-band response expected from a symmetric impedance profile (or to a high-pass spiking response if the impedance profile would have been skewed to higher frequencies). In sum, spiking resonance can be generated by the same mechanisms that generate subthreshold resonance. However, this is restricted to low input amplitudes, and therefore subthreshold resonance does not necessarily imply spiking resonance.

**Figure 4.**
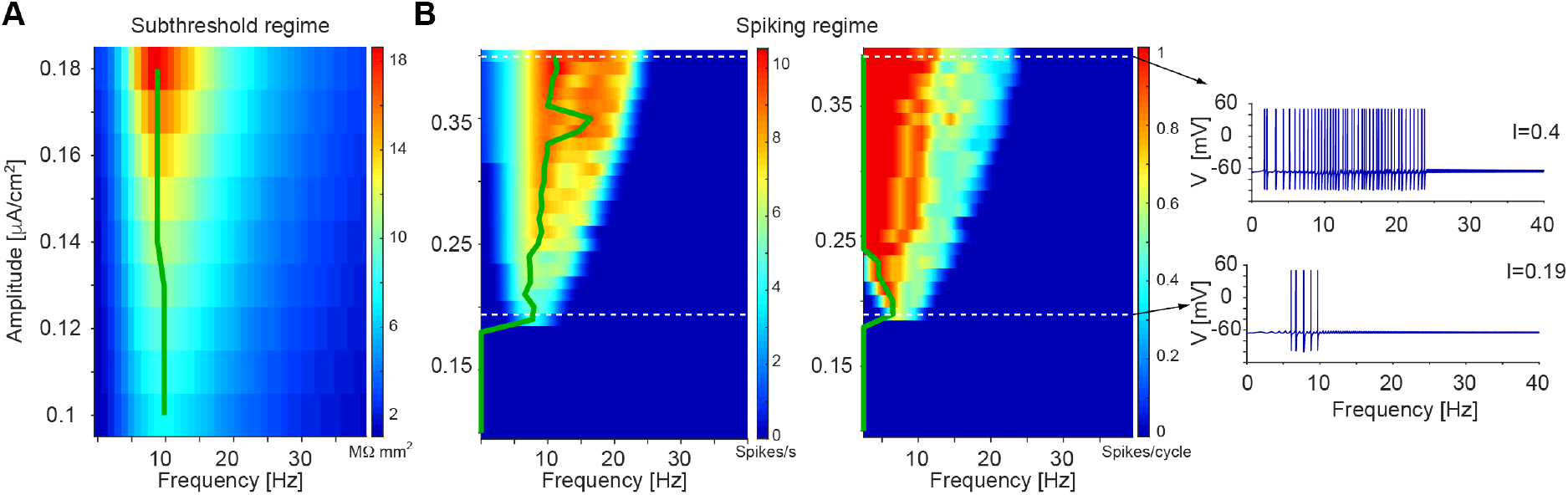
Subthreshold resonance does not necessarily imply spiking resonance. (**A**) A conductance-based spiking model neuron with “resonant” currents (same as in **Fig. 3B**) displays sub-threshold resonance with an impedance profile (heat map) that depends on input amplitude. Green line indicates the resonant (peak) frequency at each input magnitude. (**B**) When input amplitude is increased, the resonant frequency is maintained; this is captured by the green line in the left heat map which shows firing rate. However, at higher input amplitude, spikes occur at additional frequencies. This is captured by the right heat map, which shows the number of spikes per cycle. Thus, the model neuron exhibits spiking resonance at a very narrow input regime, above which spikes occur at additional frequencies. With increasing input amplitude, the subthreshold BPF is converted into a spiking LPF and thus the subthreshold resonance is not fully communicated to the spiking regime.

The second direction that we examined is whether spiking resonance may be generated without any subthreshold resonance. Conceptually, a subthreshold LPF generated by the passive (RC) properties of the membrane could interact with a spiking-domain HPF to generate spiking domain resonance. To test this idea, we employed a model that exhibits a LPF response in the subthreshold domain (**Fig. 2A**). Specifically, we designed a modified version of a leaky integrate and fire (LIF) model that includes spike-dependent calcium dynamics (**Fig. 5**). By construction, the calcium current activates only in the presence of spikes. To prevent the involvement of any currents involved in spike generation that may also generate subthreshold resonance, a spike is said to occur whenever the voltage crosses a fixed threshold (as in the standard LIF model). Without the calcium current, the model exhibited only a low-pass filter response in the subthreshold domain, and the spiking response exhibited a similar profile (**Fig. 5A**). Adding the spike-dependent calcium dynamics did not change the subthreshold response (**Fig. 5B**), but a spiking band-pass filter emerged (**Fig. 5C**). This occurred since a spike is followed by a calcium transient: a rapid increase and slower decrease of calcium, which is the basis of calcium imaging (Grienberger and Konnerth, 2012). During the calcium transient, the membrane potential was more depolarized, allowing the generation of a spike in response to a lower current input, effectively reducing spiking threshold. Thus, the occurrence of one spike favored the occurrence of another spike during a specific time window dictated mainly by the calcium activation and deactivation time constants, yielding a spiking-domain high-pass filter. Together with the subthreshold LPF, spiking resonance emerged (**Fig. 5C**). The positive feedback underlying this spiking domain resonance mechanism requires an initiator in the form of a “first” spike, which can be produced by random fluctuations. In summary, the modified LIF model with spike-dependent calcium dynamics shows that spiking resonance can be generated in the lack of any mechanisms that generate subthreshold resonance. Together, these case studies (**Fig. 4** and **Fig. 5**) clarify that independent mechanisms can generate resonance at the subthreshold and the spiking regimes.

**Figure 5.**
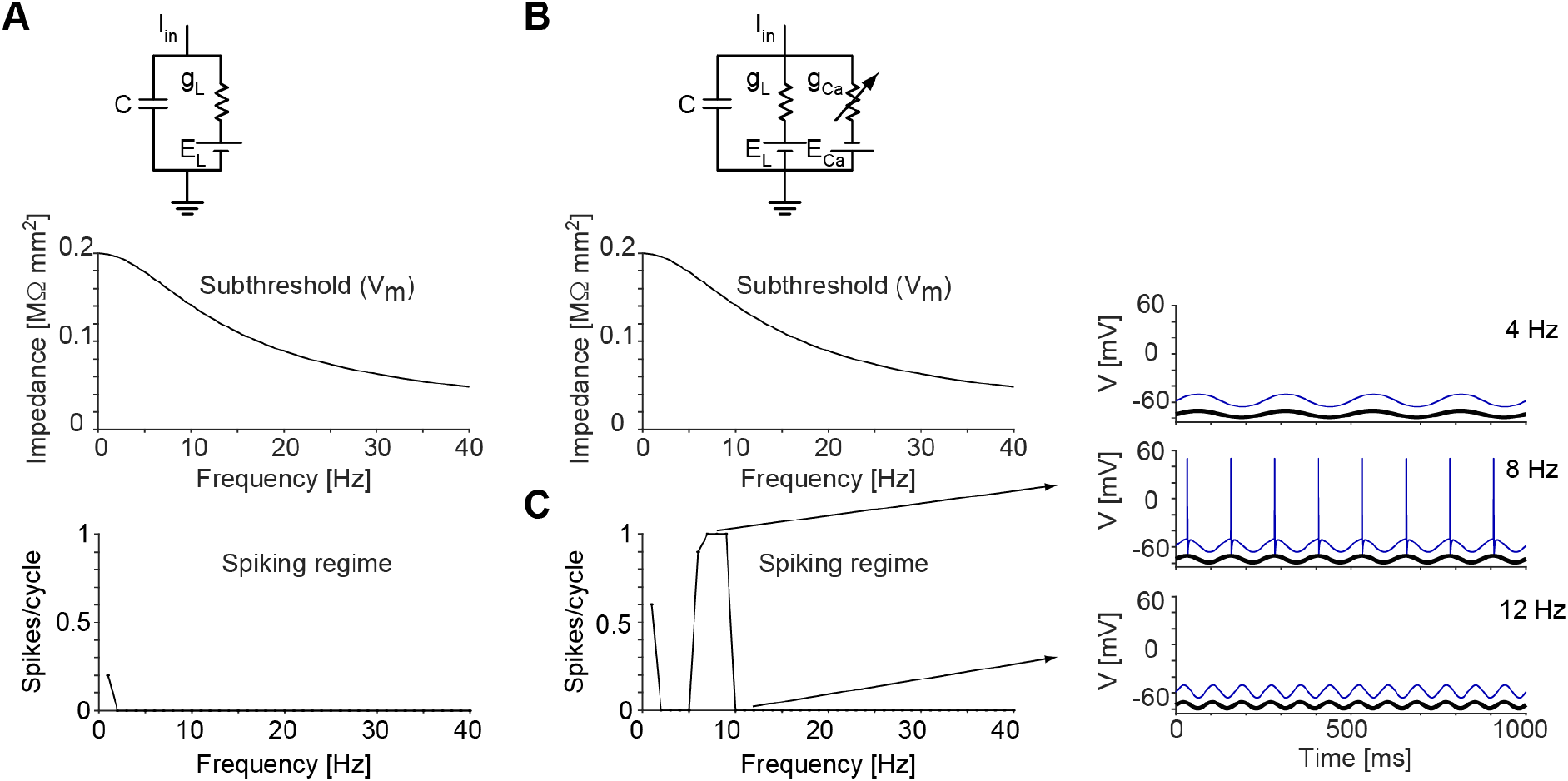
Spiking regime resonance can occur in the absence of subthreshold resonance. (**A**) A leaky integrate- and-fire (LIF) model neuron, which displays a low-pass filter (LPF) response in the subthreshold domain, also displays a LPF spiking response. (**B**) A modified LIF neuron with spike dependent calcium dynamics, yet without any inductive currents, also displays subthreshold LPF response to oscillatory input. (**C**) However, the same model displays a BPF response in the spiking regime, i.e. spiking resonance. Thus, spiking resonance is not inherited from the subthreshold regime.

### Synaptic resonance can occur independently of subthreshold or spiking resonance

Synaptic transmission displays time- and frequency-dependent properties. When presynaptic spikes form a train of a fixed frequency, the post-synaptic effect of each individual presynaptic spike may differ (Thomson et al., 1993). A decrease in the amplitude of the post-synaptic potential (PSP) between sequential presynaptic spikes reflects the presence of synaptic depression, whereas an increase of PSP amplitude between sequential spikes reflects the presence of synaptic facilitation (Zucker, 1989; Zucker and Regehr, 2002). In extracellular recordings, synaptic depression and facilitation may be quantified by comparing peak height of cross-correlation histograms triggered on isolated or sequential spikes (Fujisawa et al., 2008). To estimate frequency modulation of synaptic transmission, PSP amplitude can be measured at the end of a periodic presynaptic spike train of a given frequency, once the EPSPs have reached steady state, and compared across trains of different frequencies. In this case, synaptic depression (facilitation) has been observed as a decrease (increase) in steady-state PSP amplitude with increasing frequency (Markram et al., 1998). Synaptic resonance can then be defined as a peak in the steady-state EPSP amplitude at an intermediate input frequency (Izhikevich et al., 2003). However, it is not immediately clear if and how depression (facilitation) in the time domain translates to depression (facilitation) in the frequency domain and vice versa.

To study the interplay between the time- and frequency-dependent properties of synaptic transmission, we employed a passive membrane model of a post-synaptic cell receiving an excitatory input with time-dependent synaptic facilitation and depression (**Fig. 6A**). In this model, when the input presynaptic spike train had a low (e.g. 1 Hz) frequency, EPSP amplitude increased over time (**Fig. 6B, left**), reflecting the effect of synaptic facilitation. At high input frequencies (e.g. 100 Hz), EPSP amplitude initially increased and then decreased yielding a transient peak, reflecting the combined effect of synaptic facilitation and depression (**Fig. 6B, right**). An increase of EPSP amplitude over time was also apparent at intermediate frequencies (e.g. 10 Hz), where the steady-state EPSP amplitude had the largest values (**Fig. 6B, center**). This configuration of steady-state EPSP values corresponds to synaptic resonance (**Fig. 6D**). Thus, the time-dependent properties of synaptic transmission can give rise to (steady-state) frequency-dependent modulation of EPSP amplitude and, in particular, to synaptic resonance. This phenomenon is inherently synaptic, since it occurs without any frequencydependent modulation of the input spike train and does not require the post-synaptic cell to exhibit subthreshold or spiking resonance.

**Figure 6.**
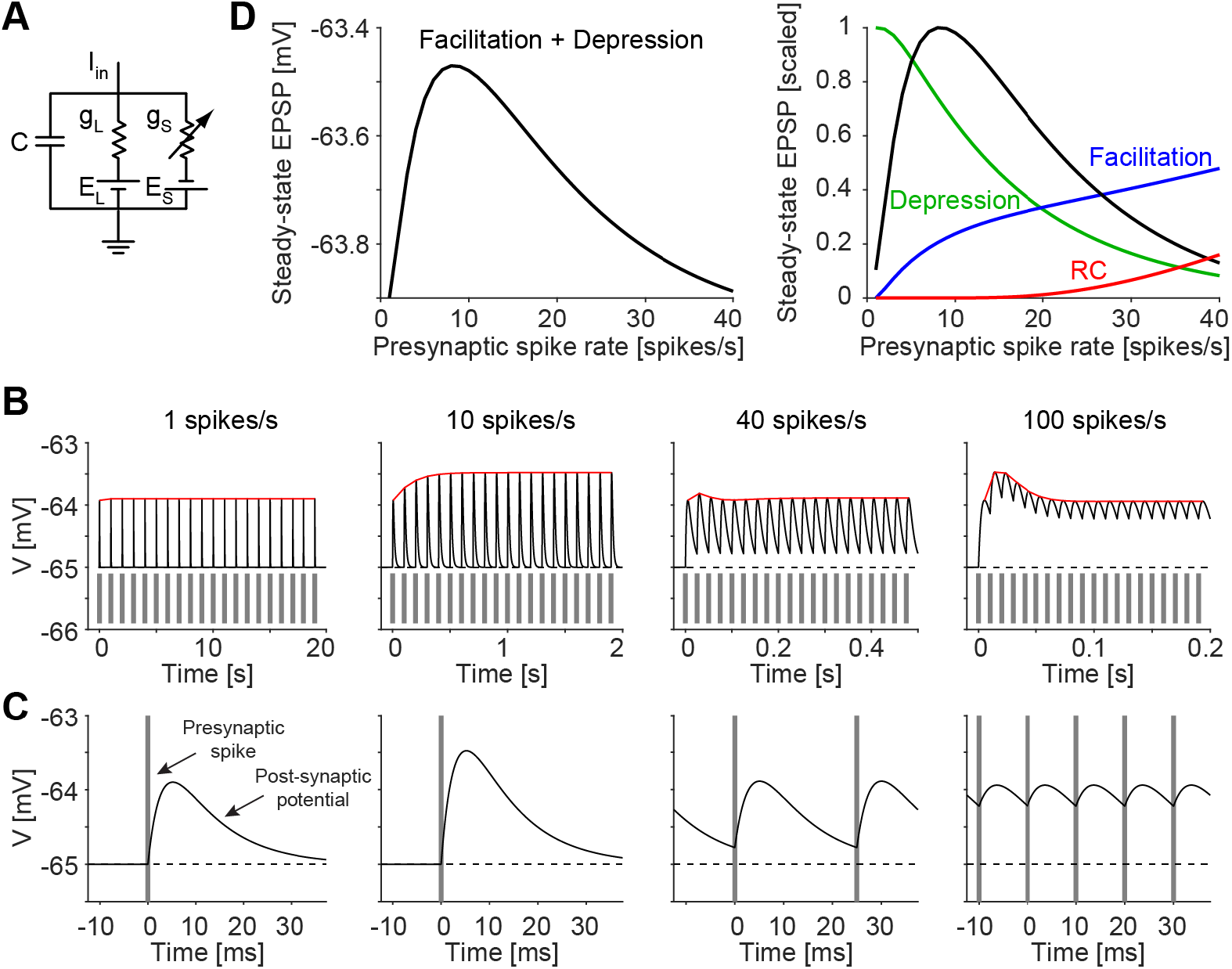
Synaptic transmission can display resonance in the lack of subthreshold or spiking resonance. (**A**) Passive membrane model with synaptic facilitation and depression. In the model, the effective synaptic conductance gs changes after every spike in a time- and history-dependent manner. (**B**) The peak amplitude of an EPSP depends on the instantaneous presynaptic firing rate. Individual membrane potential traces during input (presynaptic) spiking for the resonant model at four different rates. Input includes 20 equally-spaced spikes per firing rate. Vertical grey lines show presynaptic spike times, horizontal dashed lines show resting potential of the post-synaptic cell, and red lines trace the EPSP peaks. Note that the transient elevation and subsequent decrease at 100 spikes/s resembles a resonant curve but this is a transient response. The peak EPSP amplitude occurs at a non-zero frequency, indicating synaptic resonance. (**C**) Expanded traces at individual input rates. Vertical grey lines show spike times. (**D**) Summary curves for steady-state EPSP amplitude during synaptic resonance, synaptic depression, synaptic facilitation, and synaptic response of a passive membrane. Note that temporal summation alone, due to the RC properties of the passive membrane, introduces a high-pass filter (HPF; red trace). This property is accentuated by synaptic facilitation (blue). Synaptic depression generates a LPF (green trace), and their combination endows the EPSP with a resonant response (black trace).

### Network resonance requires frequency modulation at multiple levels of organization

Resonance can also be observed at the network level. Previously, a depolarizing oscillatory input of various frequencies (0-40 Hz) was applied optically to PV cells in freely-moving mice. Under these conditions, many PV cells responded by spiking on multiple cycles, yielding wide-band spiking with no resonant peak in the theta frequency band (Stark et al., 2013). Spiking of the putative post-synaptic pyramidal cells (PYR) was suppressed at every frequency, except at intermediate frequencies (6-10 Hz), during which spiking was induced. This phenomenon was abolished by I_h_ blockade and was termed inhibition-induced network resonance.

To determine how network resonance can be generated, we initially employed a model consisting of a single PYR. Specifically, we used a conductance-based model of the Hodgkin-Huxley type with spiking currents (same as in **Fig. 3B**). Instead of varying the amplitude of the oscillatory input current, we fixed the amplitude to a level that induced spiking resonance (Ain = 0.2 μA/cm^2^; **Fig. 4**) and varied the DC level of that current. When the current was depolarizing, spiking resonance was induced (**Fig. 7A**), consistent with the subthreshold response (**Fig. 4**). However, when the current was hyperpolarizing, no spikes were induced (**Fig. 7A**). Thus, mild oscillatory inhibition alone cannot induce spiking resonance in an isolated model cell.

**Figure 7.**
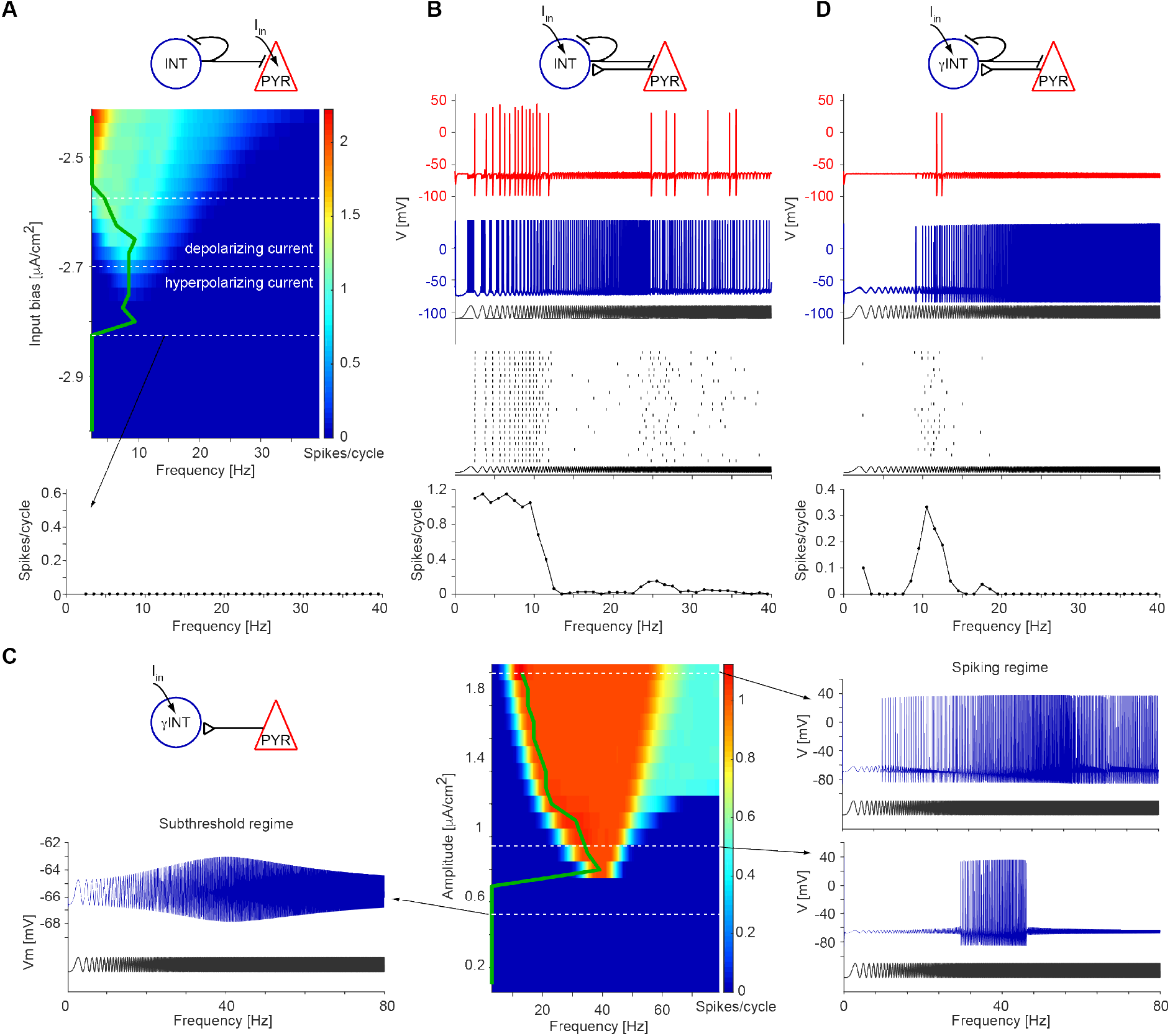
Network resonance requires frequency modulation at multiple other levels of organization. In this figure, two cells are considered: an inhibitory interneuron (INT), representing a PV-cell; and a pyramidal cell. Both cells are modeled using conductance-based models of the Hodgkin-Huxley type with spiking currents. The PYR is modeled as in **Fig. 3B**, containing *I_h_*. The INT makes a fast GABA_A_ synapse on the PYR (and on itself). (**A**) In the PYR model, currents are considered depolarizing/hyperpolarizing current if the bias is above/below −2.7 μA/cm^2^. Direct injection of hyperpolarizing sinusoidal current to the PYR at an amplitude that induces spiking resonance (0.2 μA/cm^2^; **Fig. 4**) does not induce spiking. Thus, despite the ability of the model to generate subthreshold and spiking resonance, no spiking resonance is induced in this setting. (**B**) Routing inhibition via the INT induces rebound spiking in the PYR during inhibition troughs. Here, a depolarizing current is injected to the INT, which spikes at almost every cycle. The PYR is induced to spike preferentially during the slower frequencies, and the induced spiking follows a low-pass filter pattern. No resonance is induced. (**C**) The INT model is modified by adding an M-current (*I_M_*). This endows the INT with subthreshold resonance in the gamma band (left), which is transmitted to the spiking domain (right). (**D**) The INT in the two-cell model is modified to exhibit gamma-band resonance (as in **Fig. 7C**). Considered in the 0-40 Hz range, this is a high-pass filter, which interacts with the rebound-induced low-pass filter of the PYR (**Fig. 7B**) to generate rebound spikes limited to the theta band.

Next, we used a two-cell model consisting of an interneuron (INT, representing a PV cell) that received a depolarizing oscillatory input current, connected via an inhibitory synapse to a PYR (**Fig. 7B**). The PYR was modelled as in **Fig. 7A**, whereas the INT had slightly faster dynamics than the PYR (Wang and Buzsáki, 1996) and did not have an h-current. When a chirp input was applied to the INT, that cell spiked at a wide range of frequencies, and the post-synaptic PYR was induced to spike predominantly at lower input frequencies (**Fig. 7B**). On each and every cycle of these frequencies, the PYR spikes were generated immediately after INT spikes ceased, consistent with post-inhibitory rebound spiking (Llinas and Jahnsen, 1982; Kandel and Spencer, 1961; Buhl et al., 1994; Cobb et al., 1995; Stark et al., 2013). Thus, a two-cell model in which the PYR has I_h_ and is driven by IPSPs from a periodically-activated INT could generate PYR spikes. However, the spiking response displayed only a low-pass filter response and no spiking resonance was exhibited.

To obtain spiking resonance in the network, the spiking low-pass filter of the PYR in the simple two-cell model (**Fig. 7B**) has to interact with a high-pass filter. One way to generate a HPF in this model is to include synaptic depression (Stark et al., 2013); depression rather than facilitation (**Fig. 6D**) is required since the high-pass filter is to be applied to the rebound PYR spikes. Another possibility is to extend the network, including a third cell (e.g. an OLM cell; Stark et al., 2013). Here, we present a third independent idea, namely to use a spiking high-pass filter in the INT, as observed for some of the fastspiking PV cells (**Fig. 3A**). To model a HPF in the INT, we added a slow potassium current (IM-like; Brown and Adams, 1980) to the INT. This modification endowed the INT with subthreshold resonance in the gamma band, that was transmitted to the spiking domain of the INT at intermediate input current amplitudes (**Fig. 7C**). When the INT in the two-cell model was endowed with gamma resonance, the inhibition-induced PYR spikes were limited to the theta band (**Fig. 7D**). This is due to the interaction of the low-pass filtering properties of the INT-PYR system (**Fig. 7B**) with the high-pass properties of the gamma-resonant INT (**Fig. 7C**). Thus, in this model, network resonance requires the interaction of frequency-dependent modulation at multiple levels of organization.

### Spiking phasonance can be generated in the lack of spiking resonance

Whereas resonance is defined as a peak in the response of a system to a periodic input at a non-zero input frequency, there is no constraint on the input-output phase difference. We define phasonance when the input and output exhibit zero-lag phase synchronization at a non-zero input frequency (Rotstein, 2017b). Subthreshold phasonance can occur in a model that exhibits subthreshold resonance (passive membrane with h-current; **Fig. 2C-D**). In this model, the phase of the input and output coincided at a phasonant frequency of 6 Hz, whereas the voltage amplitude was maximal at a resonant frequency of 8 Hz (**Fig. 8A**). Thus, even when both phasonance and resonance occur, the phasonant and resonant frequencies are not identical.

**Figure 8.**
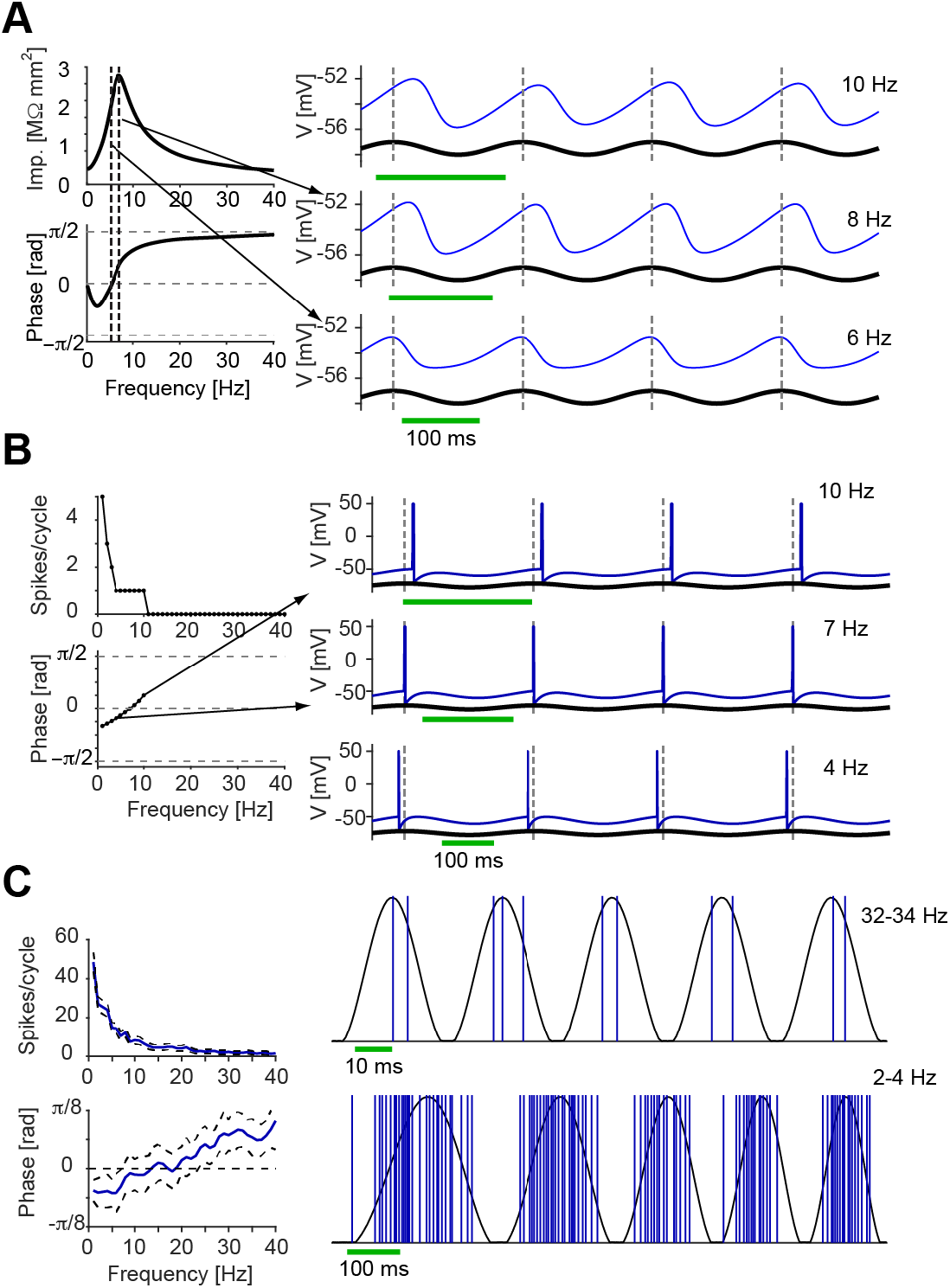
Phasonance can occur without resonance. (**A**) Subthreshold phasonance and resonance can occur at different frequencies. Phasonance, defined as a zero-lag phase difference between the input and output (input-output phase synchronization), can occur in a system exhibiting resonance. Traces show injected current and induced voltage in a leaky membrane model with *I_h_* and *I_Na,p_* (same as in **Fig. 2D**). Note that in this model, subthreshold phasonance and resonance occur at different frequencies. The frequency with zero-lag phase difference (the phasonant frequency) is 6 Hz, whereas the frequency with the maximal voltage response (the resonant frequency) is 8 Hz. (**B**) Spiking phasonance without spiking resonance in a model neuron. A LIF model (i.e. without active ionic currents) behaves as a LPF, producing spikes at a range of input frequencies without resonance. However, spike phase varies with the frequency. Spikes occur at the peak of the input current at a phasonant frequency of 7 Hz. (**C**) Spiking phasonance without spiking resonance in a CA1 interneuron. PV interneuron spiking recorded extracellularly during chirp-pattern optical activation in a freely-moving PV::ChR2 mouse. This cell does not exhibit any spiking resonance. However, spike phase is earlier for lower input frequencies, with a phasonant frequency of about 10 Hz.

For a 2D linear neuronal model, subthreshold phasonance cannot occur without subthreshold resonance (Rotstein and Nadim, 2014). For spiking models with 2D subthreshold dynamics and low input amplitude, spiking resonance can occur with no spiking phasonance (Rotstein, 2017bc). To determine whether spiking phasonance requires resonance, we used a LIF model neuron that responds to periodic inputs with low-pass spiking (**Fig. 4Ba**). Thus, the number of spikes fired at each cycle decreased monotonically with input frequency. However, for higher input frequencies the spikes occurred at later phases, crossing zero at about 7 Hz (**Fig. 8B**). In this model, spikes occurred at a fixed threshold, and phasonance was a direct outcome of the fact that the fixed time delay induced by the passive membrane corresponded to larger phase shifts at increasing input frequencies. Thus, zero-phase synchronization of the spiking output with the input current (spiking phasonance) can occur in the lack of spiking resonance.

A similar phenomenon was observed in extracellular recordings from freely-moving PV::ChR2 mice in which the PV cells were driven by a linear chirp input. The example PV cell responded with spiking at all input frequencies (up to 40 Hz), with a decreasing number of spikes per cycle (**Fig. 8C, top left**). Thus, no spiking resonance was displayed by this unit. However, the phase of the induced spikes increased when input frequency was increased (**Fig. 8C, bottom left**). While at low input frequencies, the mean spike phase was negative representing a phase advance, the mean spike phase at higher frequencies was positive, representing a phase lag (**Fig. 8C**). The phasonant frequency was higher than in the LIF model: the output spikes exhibited zero-lag phase synchronization with the input at ~10 Hz. While the mechanisms that induce phasonance in this setting may differ from those in the LIF model, the neurophysiological data indicate that neuronal spiking phasonance can occur in the lack of resonance.

## DISCUSSION

In this work, we tested the hypothesis that resonance at one level of neuronal organization is necessarily inherited from resonance at another level. From electric circuit theory it is clear that one can construct a macro-circuit consisting of two circuit components, each being able to produce resonance on its own. However, neuronal networks are naturally evolved electric circuits and it is not clear whether they have this property, primarily since the positive and negative feedback effects have different biological substrates at different levels of organization (e.g., resonant and amplifying ionic currents, excitation and inhibition, synaptic depression and facilitation). Examining four levels of neuronal organization, we found that resonance can be generated independently at each level. By combining in vivo experiments, numerical simulations of biophysically plausible models, and results from the literature, we showed that it is possible for a given system to display resonance at only one level of organization – subthreshold, synaptic, or spiking – but not in any other. In computer models, subthreshold resonance was not necessarily accompanied by spiking resonance, and spiking resonance was not necessarily accompanied by subthreshold resonance. Synaptic resonance was not necessarily accompanied by neither subthreshold nor spiking resonance (Markram et al., 1998 PNAS; Izhikevich et al., 2003). In both computer models and freely-moving mice, phasonance could occur in the lack of resonance. Thus, the mechanisms that can generate neuronal resonance at different levels of organization are distinct (**Fig. 9**). A direct implication of these observations is that when a system presents resonance and/or phasonance at multiple levels of organization, these can be derived from independent mechanisms.

**Figure 9.**
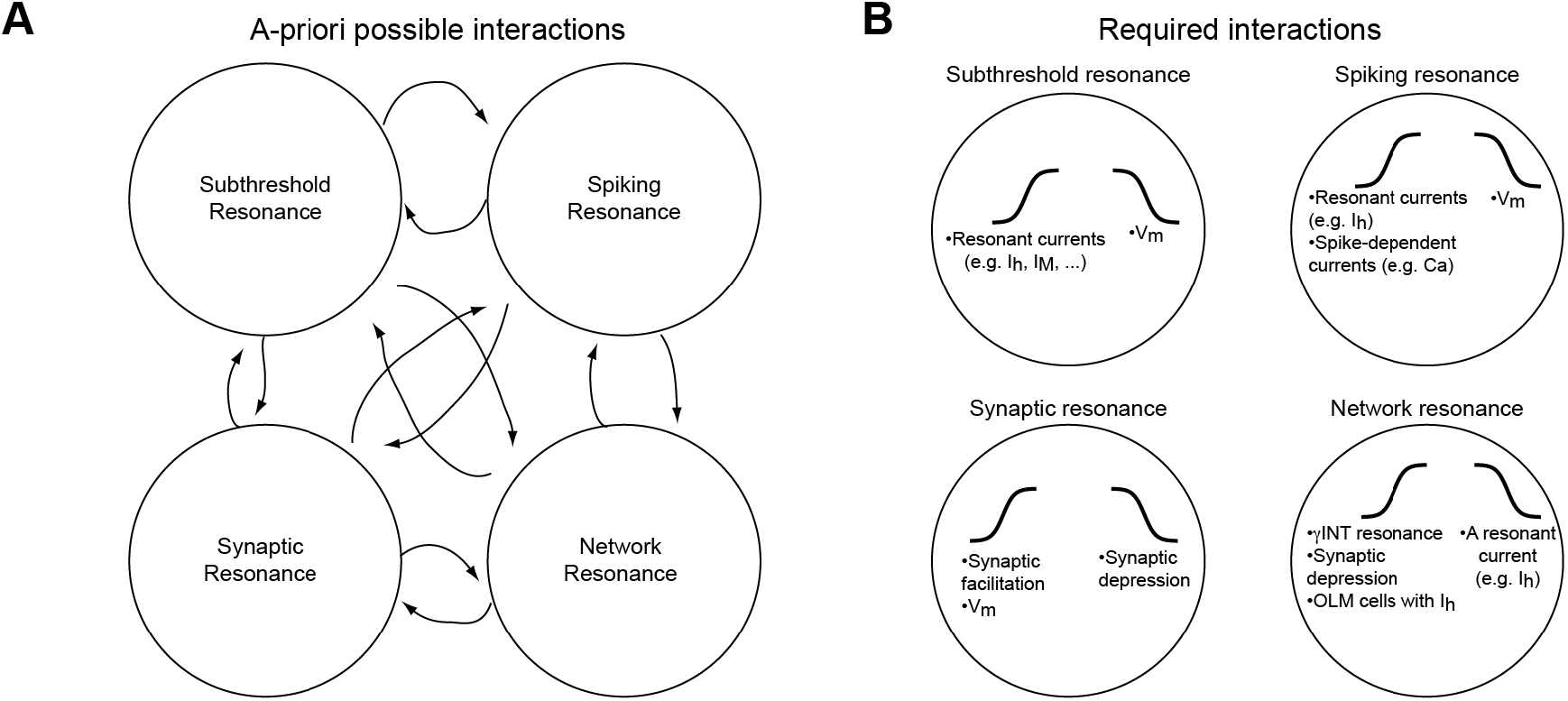
Neuronal resonance can occur independently at multiple levels of organization. (**A**) Resonance at one level may or may not be directly inherited from resonance at other levels. There are many potential mutual dependencies and many possible interactions. (**B**) However, only a handful of interactions actually take place and none are necessary. The cartoons inside each circle denote the mechanisms for neuronal low- and high-pass filters discussed in this work.

Since resonance is defined as maximal amplification of the response of a system to a periodic input at a finite non-zero input frequency band, it can be obtained by applying a combination of low- and high-pass filtering (LPF and HPF, respectively) on the input. There are at least two possible ways to generate a resonant response at a given level of organization: by using an LPF and a HPF at the same level of organization, or at different levels. In the case of subthreshold resonance, we used subthreshold LPF (passive membrane) and HPF (I_h_). The same approach was used for synaptic resonance, in which both the LPF (synaptic depression) and the HPF (facilitation) belonged to the same level of organization. However, for generating spiking resonance independently of resonance at any other level, we used a mixed approach: while the HPF was spike-dependent (calcium dynamics), the LPF was inherited from the subthreshold domain (passive membrane). A similar approach could be used for generating synaptic resonance: the temporal integration properties of a passive membrane alone induce HPF behavior (**Fig. 6D, red**) that can be combined with synaptic depression to generate synaptic resonance. Yet in all three domains (subthreshold, spiking, and synaptic), resonance could be generated without resonance at any other domain.

Inhibition-induced network resonance requires that I_h_-mediated rebound spiking in pyramidal cells (Cobb et al., 1995; Stark et al., 2013) interacts with some form of high-pass filter. Previously, depression of the inhibitory synapses (on the PYR) and interaction with a third type of cell (OLM cell) were suggested as high-pass filters (Stark et al., 2013). Here, we showed that in intact mice, PV cells depolarized with a linear chirp pattern exhibit gamma resonance, extending previous in vitro observations (Pike et al., 2000). This observation suggested a third potential mechanism for the generation of theta-band network resonance, namely the interaction of the low-pass rebound spiking filter in the PYR with a gamma-band high-pass filter in the INT (Rotstein, Ito, and Stark, 2017, SFN Abstract). Network resonance was generated without synaptic or spiking resonance in the cells that resonated in the full model. Thus, in the three models suggested so far for inhibition-induced network resonance, frequency-modulating mechanisms at multiple levels of organization are required.

The definition of resonance requires maximal amplification at the resonant frequency. Technically, maximal amplification is well-defined as a global peak in the response curve. However, in previous work, the definition of spiking domain resonance included an additional requirement, namely gain above one (Stark et al., 2013). This was motivated by the idea that minimal suppression at a given frequency band is conceptually distinct from maximal amplification. According to this concept, resonance requires amplification relative to a baseline. Yet in quantifying resonance at non-spiking domains, the gain requirement was not applied. The reason gain was not considered in analyzing subthreshold resonance is that typically the system is considered to be in steady-state (DC input without any noise) and in particular, without any stationary oscillations. How can gain be included in the definitions of subthreshold or synaptic resonance? One option is to divide the frequency-response curve by its value at DC: the impedance at zero (the resistance) for subthreshold resonance (Hutcheon et al., 1996; Boehlen et al., 2011); and the amplitude of an isolated EPSP for synaptic resonance. However, this would only scale the curves and would not compare to a baseline. A baseline for quantifying the gain of subthreshold responses could be the membrane potential variance, making the resonant effect depend on the noise in the system.

An important experimental finding is that of in vivo gamma resonance in PV cells (**Fig. 3A**). Gamma resonance in this cell type has been observed in vitro in some studies (Pike et al., 2000) but not in others (Zemankovics et al., 2010). Intrinsic oscillations were observed at gamma frequencies in PV cells (Kang et al., 2018) and at theta frequencies in other interneuron subtypes including OLM cells (Boehlen et al. 2011; Chapman and Lacaille, 1999; Zemankovics et al., 2010). To our knowledge, this is the first report of gamma resonance in intact animals. However, the underlying mechanisms were not determined. First, since synaptic blockers were not used in the present experiments, we cannot absolutely discard the possibility that the generation of gamma resonance in PV cells itself has a network component. Second, even if the mechanism is intrinsic to the PV cells themselves, the ionic currents involved were not determined. In our gamma resonance INT model, we used potassium currents with faster dynamics than the standard M-type slow potassium current, but slower than the spiking transient potassium current. This slow potassium current was an educated guess, based on the type of interactions that produce resonance in other cell types, and the fact that h-currents have not been found to be prominent in fast spiking interneurons. However, it is plausible that other types of currents (e.g., A-type potassium or calcium) are responsible for the in vivo gamma resonance observed.

Another finding of conceptual importance is that of in vivo non-resonating oscillations (**Fig. 1**). On the basis of computational and in vitro experimental evidence, resonance and intrinsic oscillations have been considered to be tightly related (Lamp and Yarom, 1997; Hutcheon and Yarom, 2000).

However, theoretical studies showed that intrinsic oscillations and resonance can occur independently of each other (Richardson et al., 2003; Rotstein and Nadim, 2014), establishing that resonance is not merely uncovering the ability of a system to oscillate. To our knowledge, this is the first experimental report of independence between intrinsic oscillations and resonance. These results are consistent with the fact that intrinsic oscillations are a property of an unforced system (e.g., a neuron or network), which results from the intrinsic properties of the system. In contrast, resonance (and the impedance profile) is a property of the interaction between the system and the oscillatory input in the frequency domain (Rotstein, 2014; Leiser and Rotstein, 2019), in much the same way as phase and spike-time response curves are in the time domain (Pervouchine et al., 2006; Smeal et al., 2010).

We have presented several original experimental results and several novel computational models, and have rejected the hypothesis that resonance at one level of organization necessarily requires resonance at another level. While doing so, we set the infrastructure for a theoretical framework for investigating the mechanisms underlying the generation of neuronal network oscillations, taking into account the interplay of the intrinsic properties of the participating neurons, synaptic connectivity, and network topology. This framework will enable studies of neuronal networks where the interactions between periodic inputs, intrinsic currents and network effects are important (Lisman, 2005; Iaccarino et al., 2016; Helfrich et al., 2019), different networks entrain each other (Sirota et al., 2008; Fries, 2015), and/or the precise coordination between periodic input and spiking output are enhanced or disrupted (Bi and Poo, 2001; Lakatos et al., 2008; Vierling-Claassen et al. 2008).

## ACKNOWLEDGEMENTS

This work was funded by a CRCNS grant jointly from the United States-Israel Binational Science Foundation (BSF #2015577; ES), Jerusalem, Israel, and the United States National Science Foundation (NSF #1608077; HGR); and by an ERC Starting Grant (#679253; ES). We thank Amir Levi, Hadas Sloin, Shirly Someck, and Lidor Spivak for help collecting data; to Takuya Ito for preliminary simulations; and to Roni Gattegno, Drew Headley, Takuya Ito, Amir Levi, Rodrigo Pena, Shir Sivroni, Hadas Sloin, Shirly Someck, Lidor Spivak, and Alexander Tarnavsky for insightful comments and productive discussions.

## METHODS

### Animals and surgery

Eight freely-moving mice (three CaMKII::ChR2, three PV::ChR2, one VIP::ChR2, and one CCK::ChR2) were used in this study. To express ChR2 in pyramidal cells (PYR), we injected a viral vector (rAAV5/CaMKIIa-hChR2(h134R)-mCherry; viral titer estimated at 5.6×10^12^ IU/mL; University of North Carolina viral core facility, courtesy of K. Deisseroth) stereotactically (Kopf) into neocortex and hippocampus of wild-type mice at 8 different depths (AP −1.6, ML 1.1, DV 0.4 to 1.8 at 0.2 mm increments; 50 nl/site; Nanoject III, Drummond). To express ChR2 in parvalbumin-immunoreactive (PV) cells, we cross-bred male Ai32 mice homozygous for the Rosa-CAG-LSL-ChR2(H134R)-EYFP-WPRE conditional allele (#012569, Jackson Labs) with homozygous female PV-Cre mice in which Cre recombinase expression is directed to PV expressing cells (#008069, Jackson Labs). To express ChR2 in vasoactive intestinal peptide-immunoreactive (VIP) or cholecystokinin-immunoreactive (CCK) cells, we cross-bred the Ai32 males with homozygous female VIP-Cre (#010908, Jackson Labs) or with CCK-Cre (#012706, Jackson labs) female mice.

All animals were implanted with multi-site diode-probes on movable microdrives following previously-described procedures (Stark et al., 2012). Briefly, diode-probes were constructed by coupling miniature blue LEDs (470 nm; LB P4SG, Osram) or blue laser diodes (LDs; PL450, Osram) to glass optical fibers etched to a point (AFS50/125, Thorlabs) or to polyimide fibers (WF50/60/70P, NA=0.28, Ceramoptec). Each diode-fiber assembly was attached to a single shank of a commercially-available silicon probe (NeuroNexus). Three probe types were used: (1) A1×32 Edge with 100 μm vertical separation between recording sites, used in two of the three CaMKII::ChR2 mice; (2) 32-site/4-shank probes, Buzsaki32, 20 μm vertical separation, used in the three PV::ChR2 mice; and (3) A1×32 Edge with 20 μm vertical separation between recording sites, used in the last three mice. At the time of implantation, the light output at the tip of the shanks, when driven with a 50 mA current, was 31±4 μW (mean±SEM; n=15 470 nm LEDs) and 124±40 (n=3 450 nm LDs). Fiber tips were located ~50 μm above the top recording site; this configuration yielded maximal blue light intensity of approximately 1.5 mW/mm^2^ at the target region, corresponding to the center of the shank (4-shank probes) or the top recording site (linear probes). In linear probes with 100 μm spacing, a second fiber was etched to a point and attached at a slight angle just above the 25^th^ recording site. Mice were tethered by one ultralight cable for multi-channel neuronal recordings and a second cable for multi-channel optical stimulation.

### Recording procedures and photostimulation

Neural activity was amplified, filtered, multiplexed, and digitized on the headstage (0.1–7,500 Hz, x192; 16 bit, 20 kHz; RHD2132, Intan Technologies). Animals were equipped with a 3-axis accelerometer (ADXL-335, Analog Devices) for monitoring head-movements, two head-mounted LEDs for online video tracking (Gaspar et al., 2019), or both. All shanks were equipped with diodes, each driven by a separate channel of a custom-made 16-channel current source, controlled by a programmable DSP (24.414 kHz, RX8; Tucker-Davis Technologies) via MATLAB (The MathWorks, Natick, MA). The applied currents were recorded by separate channels of the data acquisition system.

Recordings were carried out in the home cage during spontaneous behavior. After each session, the probe was either left in place or advanced in 35–70 μm steps and the brain was allowed to settle overnight. At each location in the brain, neuronal activity was inspected for spontaneous spiking activity, and if encountered, a full recording session (3-4 hours) was conducted. The session consisted of a baseline period of at least 15 min, followed by light stimulation. Initial photostimulation was performed by each diode separately using 50-70 ms pulses at the minimal light intensity that evoked an effect detectable by visual inspection during the experiment (“spiking threshold”, defined using the driving current *I_max_*) and at multiple intensities above and below that level (log-spaced multiples of *I_max_*; typically 0.5, 1, 2, and 4). Subsequently, we used either discrete sinusoids of the form

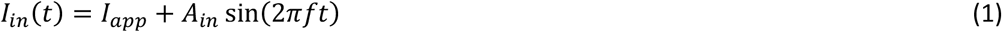

or a linear chirp (ZAP; Puil et al., 1986) of the form

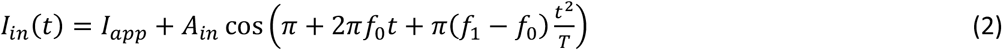

where *I_app_* is a fixed (time-independent) DC bias current, and *A_in_* is the amplitude of the time-dependent input. In the case of discrete sinusoids, input frequency *f* was typically varied from 0.1 Hz to 40 Hz at 0.25 increments. For linear chirps, we used *f_0_* = 0 Hz and *f_1_* = 40 or 100 Hz with T=10 s or 20 s, respectively. The actual currents applied to the diodes used *I_app_*=0 and *A_in_*=1, and then scaled by *I_max_* as follows. For LEDs (which exhibited linearly-increasing light output at increasing currents), the actual current was *I(t)=I_max_(I_in_(t)+1)/2*. For LDs (which exhibit linear output only in a narrow range above *I_TH_*) the applied current was *I(t)*=*I_TH_+(I_max_−I_TH_)(I_in_(t)+1)/2*.

### Spike detection, sorting, and cell type classification

For offline analysis, spike waveforms were extracted from the wide-band recorded signals and sorted into individual units (Stark et al., 2013). In multi-shank probes, waveforms were linearly detrended, projected onto a common basis obtained by principal component analysis of the data, and sorted automatically (Harris et al., 2000) followed by manual adjustment. In dense linear probes, spikes were detected and sorted using a matching-pursuit algorithm that used spike-based templatematching (Pachitariu et al., 2016), followed by manual adjustment. Only well-isolated units (amplitude >50 μV; L-ratio <0.05, Schmitzer-Torbert et al., 2005; interspike interval index <0.2, Fee et al., 1996) were used. Subsequently, each unit was tagged as excitatory/inhibitory (based on peaks/troughs in the short-time (±5 ms) pair-wise cross correlation; P < 0.001, convolution test; Stark and Abeles, 2009) and/or classified as a putative PYR or INT (based on a Gaussian-mixture model; P < 0.05; Stark et al., 2013). We recorded a total of 1155 well isolated cells from CX and CA1 of the eight mice during 60 sessions. Of these, 913 were PYR and 242 were INT.

### Models

We used biophysical (conductance-based) models, following the Hodgkin-Huxley (HH) formalism (Hodgkin and Huxley, 1952; Ermentrout and Terman, 2010). Models consisted of a set of coupled ordinary differential equations. A detailed description of the different models used is provided below.

### Numerical methods

All numerical simulations were carried out using custom code written in MATLAB (The Mathworks, Natick, MA). Numerical integration was done using the explicit second-order Runge-Kutta endpoint (modified Euler) method with integration time step *dt* ms and simulation duration of *T* ms. As current input, we used either discrete sinusoids (**equation 1**) or linear chirps (**equation 2**).

### Models for subthreshold resonance

To model subthreshold resonance (**Fig. 2**), we used a two-dimensional conductance-based model of the HH type without spiking dynamics. Thus, the only ionic currents were persistent sodium with instantaneous activation (I_Na,p_), and h-current (I_h_) with voltage-dependent dynamics. In this model, the resonance induced by I_h_ is amplified by I_Na,p_. The model equations were:

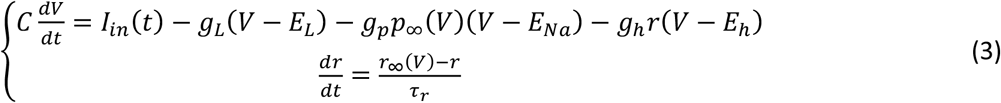

The steady-state values of the gating variables for the persistent sodium and h-current had voltagedependent sigmoid forms:

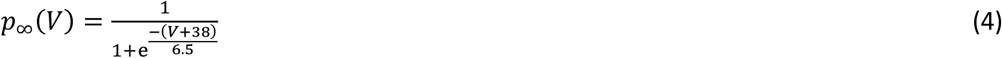

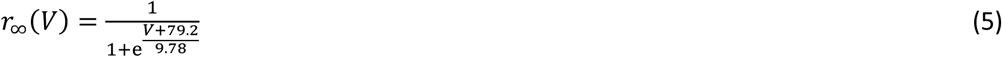

To model a passive membrane (**Fig. 2A**), we set the conductance of the persistent sodium (g_p_) and the h- (g_h_) currents to zero. To model a high-pass filter (**Fig. 2B**), we set g_p_ to zero and reduced C to 0.1. To model the basic band-pass filter (**Fig. 2C**), we set g_p_ to zero. For **Fig. 7D**, the full model was used. Other parameters used are detailed in **Table 1**.

**Table 1.** Parameters used in modeling subthreshold resonance (**Fig. 2**).

| Parameter | Value | Units |
| --- | --- | --- |
| C | 1 | $\mu\text{F}/\text{cm}^2$ |
| $g_L$ | 0.1 | $\text{mS}/\text{cm}^2$ |
| $E_L$ | -65 | mV |
| $g_p$ | 0.1 | $\text{mS}/\text{cm}^2$ |
| $E_{\text{Na}}$ | 55 | mV |
| $g_h$ | 1 | $\text{mS}/\text{cm}^2$ |
| $E_h$ | -20 | mV |
| $\tau_h$ | 100 | ms |
| $I_{\text{app}}$ | -1.8 | $\mu\text{A}/\text{cm}^2$ |
| $A_{\text{in}}$ | 0.09 | - |
| dt | 0.1 | ms |
| T | 5000 | ms |

### Models for spiking resonance

To model spiking resonance (**Fig. 3B**; **Fig. 4**; **Fig. 7A**), we used a conductance-based model of the HH type with instantaneous activation of sodium channels (Olufsen et al., 2003). In addition to the dynamics on the membrane potential (V), sodium inactivation (h), and delayed-rectifier potassium (n), the model included dynamics on the h-current gating variable (r; Poolos et al., 2002; Zemankovics et al., 2010), yielding a 4D system:

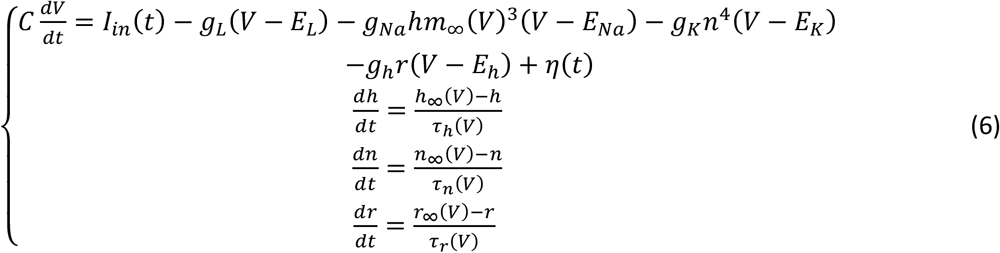

The gating variables (*x=m,n,h,r*) had voltage-dependent time constants (*τ_x_*) and steady-state values (*x_∞_*) as follows:

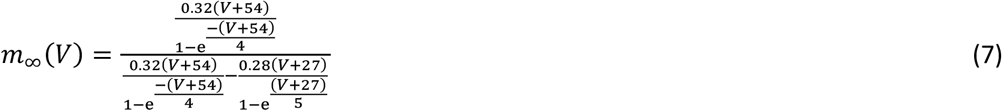

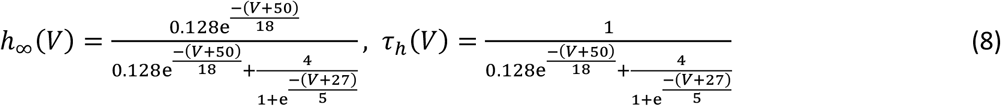

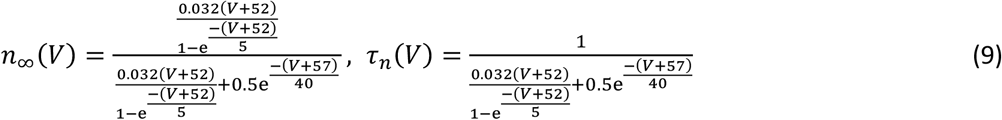

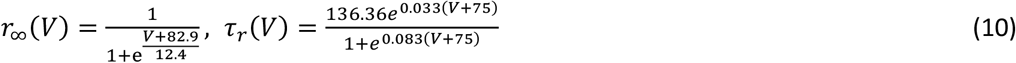

This model also included an additive noise term *η(t)* (in **equation 6.1**) that was generated by random sampling from a zero-mean Gaussian distribution with variance *σ^2^*. Other parameters used are detailed in **Table 2**.

**Table 2.** Parameters used in modeling spiking resonance (**Fig. 3** and **Fig. 4**).

| Parameter | Value | Units |
| --- | --- | --- |
| C | 1 | $\mu\text{F}/\text{cm}^2$ |
| $g_L$ | 0.1 | $\text{mS}/\text{cm}^2$ |
| $E_L$ | -67 | mV |
| $g_{\text{Na}}$ | 100 | $\text{mS}/\text{cm}^2$ |
| $E_{\text{Na}}$ | 50 | mV |
| $g_K$ | 80 | $\text{mS}/\text{cm}^2$ |
| $E_K$ | -100 | mV |
| $g_h$ | 0.485 | $\text{mS}/\text{cm}^2$ |
| $E_h$ | -33 | mV |
| $\sigma$ | 0.1 | mV |
| $I_{\text{app}}$ | -2.7 | $\mu\text{A}/\text{cm}^2$ |
| $A_{\text{in}}$ | 0.18 | - |
| dt | 0.025 | ms |
| T | 10000 | ms |

To model spiking resonance in the lack of subthreshold resonance (**Fig. 5**), we used a leaky integrate-and-fire (LIF) model, modified to include spike-dependent calcium dynamics and voltage reset. This model also included an additive noise term as described above (**equation 6**). The model equations were:

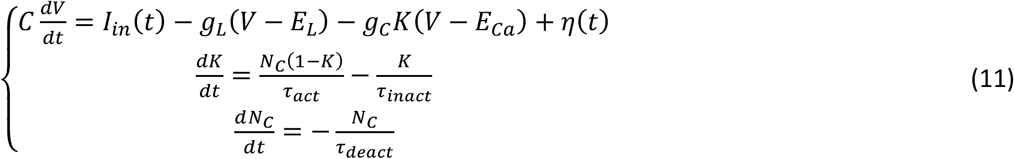

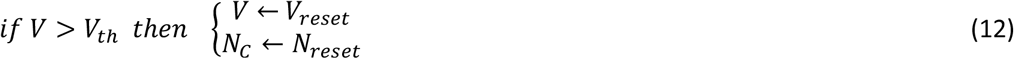

The purpose of constructing this model was to generate a spike-dependent high-pass filter in a system that has an underlying subthreshold low-pass filter. The physiological rationale is that following a spike, there is increased calcium influx, further increasing depolarization; this effectively reduces the spiking threshold to current input at the same level. Thus, at another cycle of input that occurs shortly after first spike, there will be another spike – even if the current is insufficient to generate a spike without the calcium influx. However, if the next cycle occurs later, the intracellular calcium level will have already gone back to steady-state level.

To model this, the temporal filtering properties were modeled by a gated Calcium channel. The calcium gating variable *K* is limited to the range [0,1] and represents the probability of the gate to be open. Once a spike occurs, *N_c_* is instantaneously reset to a non-zero value (*N_reset_*) and then slowly (with *τ_deact_*) decays towards zero. While *N_c_* is non-zero, the gate opens slowly (*K* is activated towards 1 with *τ_act_/N_c_*), and rapidly inactivates (decays to zero with *τ_inact_*). When activation is very fast or inactivation is very slow, calcium conductance remains high long after a spike, providing additional depolarization at multiple current input frequencies, generating spike bursts at every input cycle. When the activation is slow and inactivation is fast, *K* remains relatively high only for a short time after a spike. The parameters used (detailed in **Table 3**) favor the latter scenario.

**Table 3.**
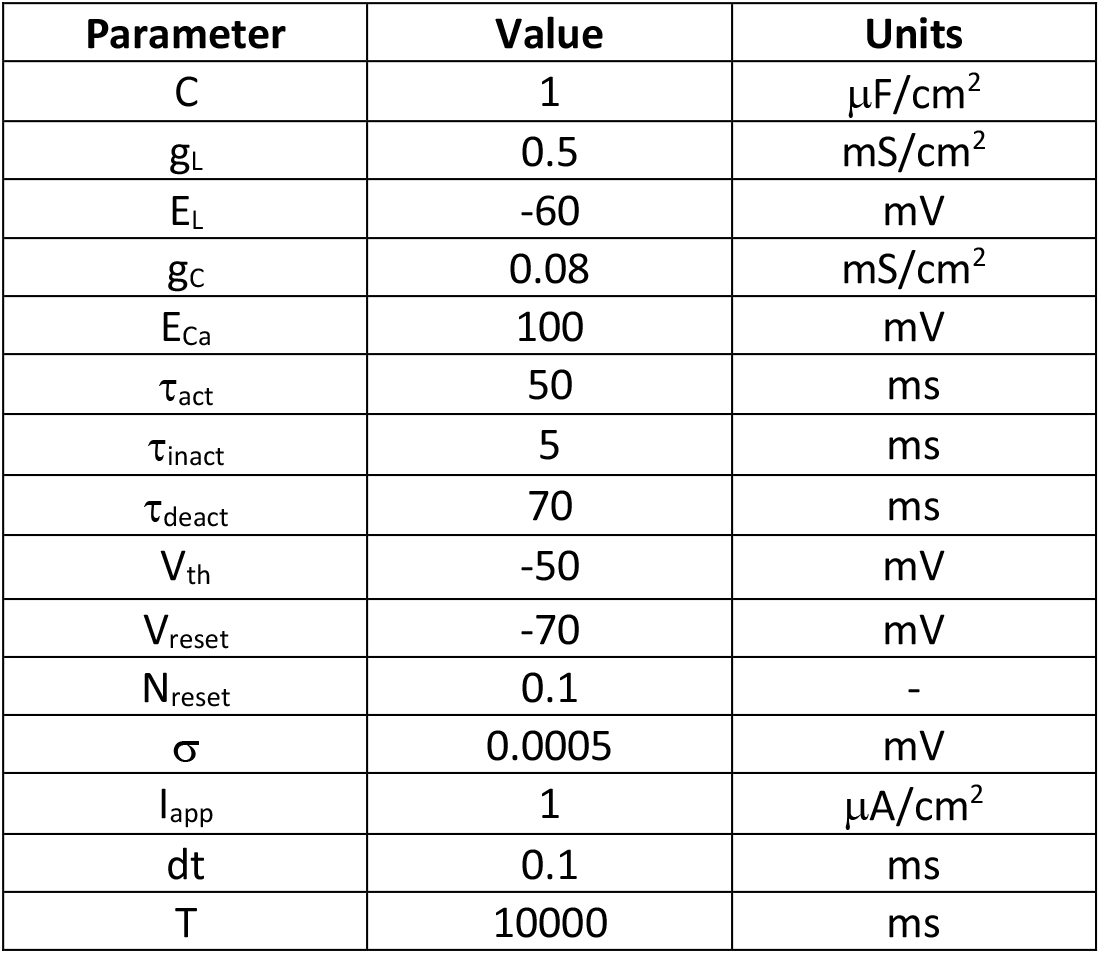
Parameters used in modeling spiking resonance without subthreshold resonance (**Fig. 5**).

**Table 4.**
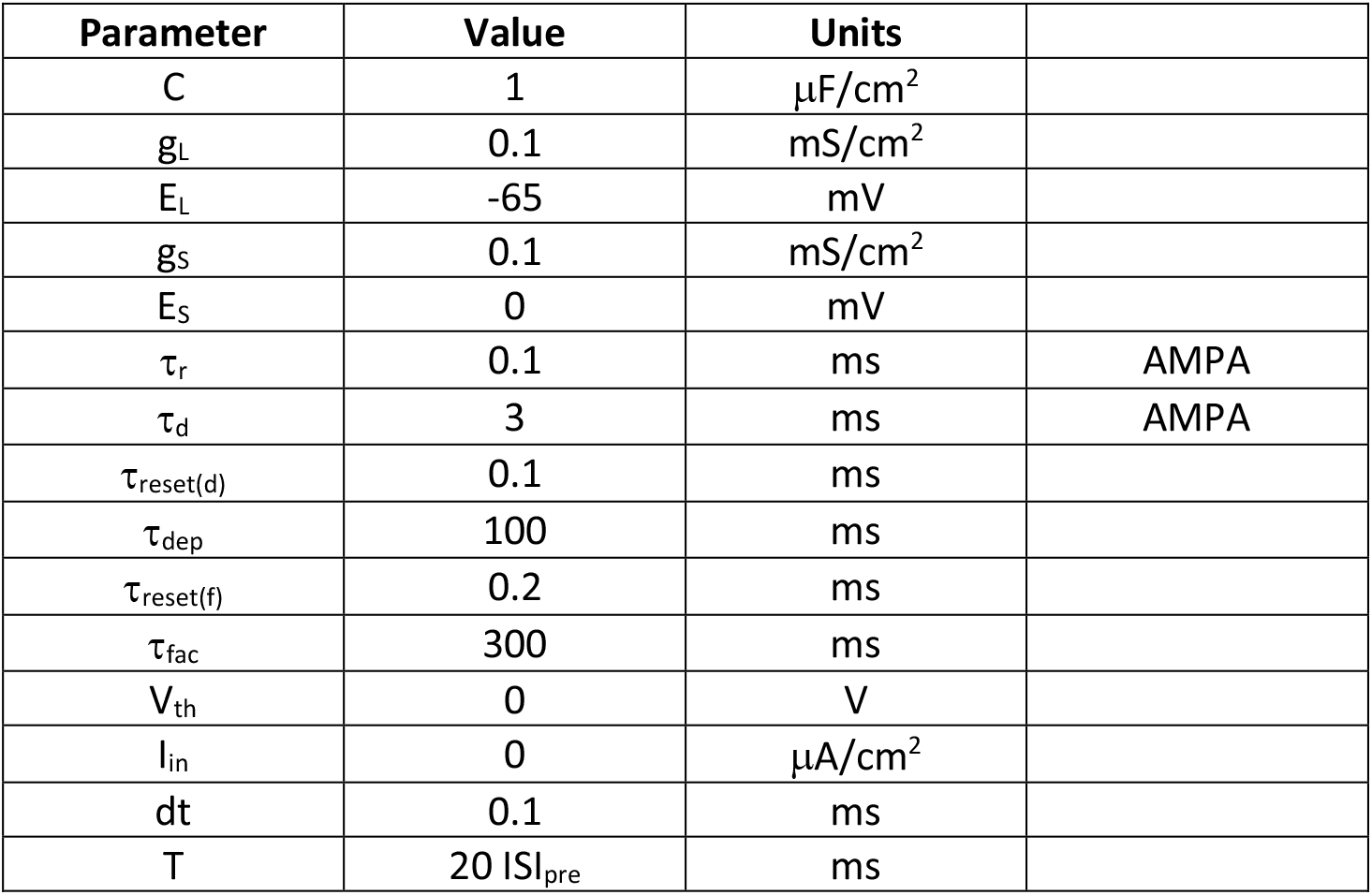
Parameters used in modeling synaptic resonance (**Fig. 6**).

| Parameter | Value | Units |  |
| --- | --- | --- | --- |
| C | 1 | $\mu\text{F}/\text{cm}^2$ | |
| $g_L$ | 0.1 | $\text{mS}/\text{cm}^2$ | |
| $E_L$ | -65 | mV | |
| $g_S$ | 0.1 | $\text{mS}/\text{cm}^2$ | |
| $E_S$ | 0 | mV | |
| $\tau_r$ | 0.1 | ms | AMPA |
| $\tau_d$ | 3 | ms | AMPA |
| $\tau_{reset(d)}$ | 0.1 | ms | |
| $\tau_{dep}$ | 100 | ms | |
| $\tau_{reset(f)}$ | 0.2 | ms | |
| $\tau_{fac}$ | 300 | ms | |
| $V_{th}$ | 0 | V | |
| $I_{in}$ | 0 | $\mu\text{A}/\text{cm}^2$ | |
| dt | 0.1 | ms |  |
| T | 20 $ISI_{pre}$ | ms | |

An integrate- and-fire type of spiking mechanism was used in this model, to ensure that any spiking resonance is not generated by the currents used to describe the spikes themselves in more detailed biophysical models (e.g. **equation 6**).

### Model for synaptic resonance

To model synaptic resonance (**Fig. 5**), we used a passive membrane model with synaptic facilitation and depression:

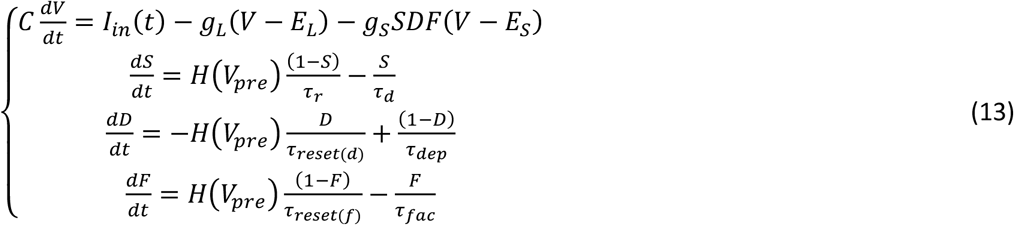

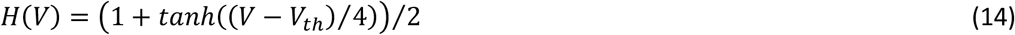

In **equations 13** and **14**, *V_pre_* and *V* represent the membrane potential of the pre- and post-synaptic neurons, respectively. The synaptic variable *S* is limited to the range [0,1] and represents the probability of the synapse to be open (or equivalently, the fraction of open synapses). During each presynaptic spike, the first term in **equation 13.2** is activated using a time-independent sigmoid function (**equation 14**). This allows the synaptic variable to grow. Once the spike is over, the first term is no longer active, and the synaptic variable decays to zero exponentially with a time constant *τ_s_* (it may not reach zero if another spike comes before this happens).

To model synaptic depression, the synaptic variable *S* is multiplied by a factor *D*, also limited to the range [0,1]. After each spike, *D* slowly recovers towards its steady state value of 1, with time constant *τ_dep_*, which determines the temporal extent of depression (**equation 13.3**). Since additional spikes may occur during recovery, the process is history-dependent. Note that the synaptic variable *S* in (**equation 13.2**) is also, in principle, history-dependent, representing synaptic summation. However, the synaptic decay time constant *τ_d_* for AMPA (and GABA_A_) is much smaller than those used for modeling depression.

To model synaptic facilitation, we introduced another multiplicative factor *F*, also limited to the [0,1] range. The dynamics of *F* follow the same principle as depression (**equation 13.4**), yet in an opposite direction: during every spike, *F* rapidly increases towards 1, and in between spikes it relaxes to zero with a slower time constant *τ_fac_*.

Both *D* and *F* affect the synaptic dynamics in a multiplicative way, namely the synaptic function *S* is multiplied by *D* and *F*. In the absence of depression or facilitation, the corresponding variable is replaced by 1. Together, the product *DF* represents the probability of presynaptic release. We note that the depression model is similar to the one proposed by Manor and Nadim (2001), and that the present model generalizes the binary rules in the models proposed by Markram et al. (1998) and Dayan and Abbot (2001).

### Models for network resonance

To model inhibition-induced network resonance (**Fig. 7D**), we used a two-cell network of conductance-based neurons of the HH type with instantaneous activation of sodium channels, consisting of a mutually connected excitatory cell (a PYR) and an INT (Borgers et al., 2012). The PYR was modeled using the 4D model described above (**equation 6**), with the voltage dynamics extended to include synaptic input from the INT. Denoting the membrane potential of the PYR by *V^e^* and the membrane potential of the INT by *V^i^*, the full model for the PYR was

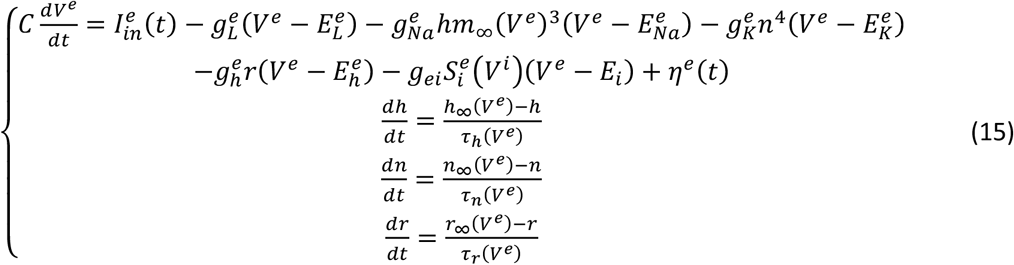

with gating variables described by **equations 7–10** and parameter values as described in **Table 5** (same as those in **Table 2**). Synaptic connections were modeled as in Ermentrout and Kopell (1998) and Borgers et al. (2012). Specifically, the inhibitory synapse on the PYR was described by a maximal conductance *g_ei_*, reversal potential *E_i_*, and a synaptic variable ranging [0,1] denoted by 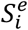. The synaptic variable was modeled using

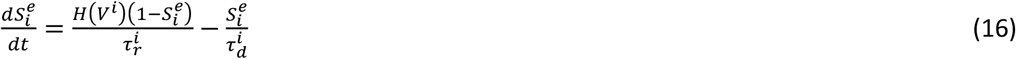

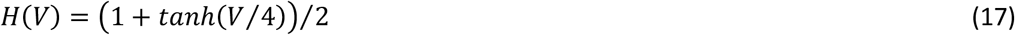

**Table 5.** Parameters used in modeling network resonance (**Fig. 7**).

| Parameter | Value | Units | Notes |
| --- | --- | --- | --- |
| $C^e$ | 1 | $\mu\text{F}/\text{cm}^2$ | |
| $g_L^e$ | 0.1 | $\text{mS}/\text{cm}^2$ | |
| $E_L^e$ | -67 | mV | |
| $g_{\text{Na}}^e$ | 100 | $\text{mS}/\text{cm}^2$ | |
| $E_{\text{Na}}^e$ | 50 | mV | |
| $g_K^e$ | 80 | $\text{mS}/\text{cm}^2$ | |
| $E_K^e$ | -100 | mV | |
| $g_h^e$ | 0.48 | $\text{mS}/\text{cm}^2$ | |
| $E_h^e$ | -33 | mV | |
| $I_{\text{app}}^e$ | -2.95 | $\mu\text{A}/\text{cm}^2$ | |
| $\sigma^e$ | 0.1 | mV | |
| $C^i$ | 1 | $\mu\text{F}/\text{cm}^2$ | |
| $g_L^i$ | 0.1 | $\text{mS}/\text{cm}^2$ | |
| $E_L^i$ | -65 | mV | |
| $g_{\text{Na}}^i$ | 35 | $\text{mS}/\text{cm}^2$ | |
| $E_{\text{Na}}^i$ | 55 | mV | |
| $g_K^i$ | 9 | $\text{mS}/\text{cm}^2$ | |
| $E_K^i$ | -90 | mV | |
| $g_M^i$ | 4 | $\text{mS}/\text{cm}^2$ | Only in **Fig. 6C, 6D** |
| $I_{\text{app}}^i$ | -0.8 | $\mu\text{A}/\text{cm}^2$ | |
| $\sigma^i$ | 0.1 | - | |
| $\tau_r^i$ | 0.3 | Ms | GABA<sub>A</sub> |
| $\tau_d^i$ | 9 | Ms | GABA<sub>A</sub> |
| $\tau_r^e$ | 0.1 | Ms | AMPA |
| $\tau_d^e$ | 3 | Ms | AMPA |
| $E_i$ | -80 | mV | |
| $E_e$ | 0 | mV | |
| $g_{ei}$ | 0.27 | $\text{mS}/\text{cm}^2$ | INT to PYR |
| $g_{ie}$ | 0.15 | $\text{mS}/\text{cm}^2$ | PYR to INT |
| $g_{ii}$ | 0.05 | $\text{mS}/\text{cm}^2$ | INT to INT |
| dt | 0.025 | ms |  |
| T | 10000 | ms |  |

Thus, whenever the presynaptic unit (the INT) spiked, the synaptic variable rose (towards 1) with time constant 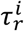 and decayed (towards zero) with time constant 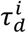; these time constants corresponded to those of fast inhibitory synapses (GABA_A_; **Table 5**).

The INT was a modeled with dynamics on the membrane potential (*V*), sodium inactivation (*h*), and delayed-rectifier potassium (*n*) (Wang and Buzsáki, 1996). To model gamma resonance in the INT, the model was extended to include a non-inactivating potassium current (*q*) with dynamics similar to but slower than an M-current (Brown and Adams, 1980). The full model also included synaptic input from the PYR and from the INT to itself, and was

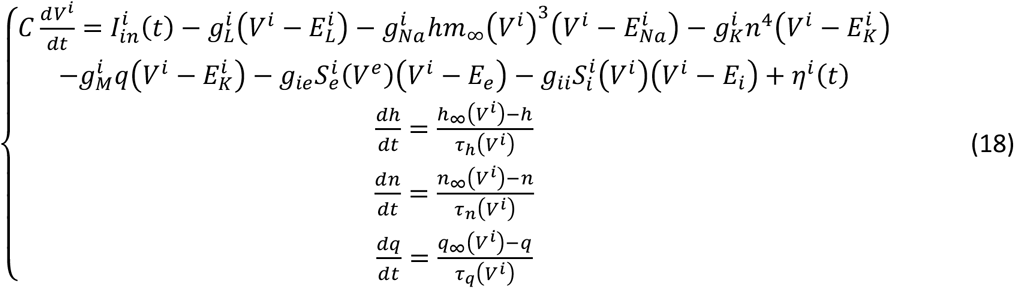

The gating variables for the INT (*x=m,n,h,q*) had voltage-dependent time constants (*τ_x_*) and steadystate values (*x_∞_*) as follows

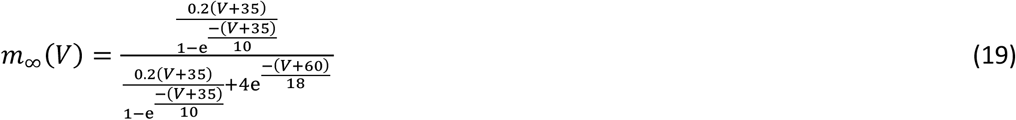

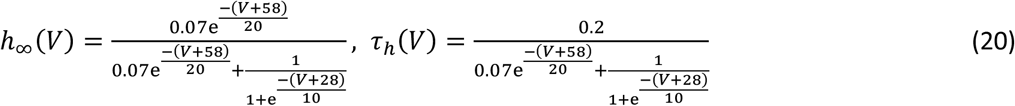

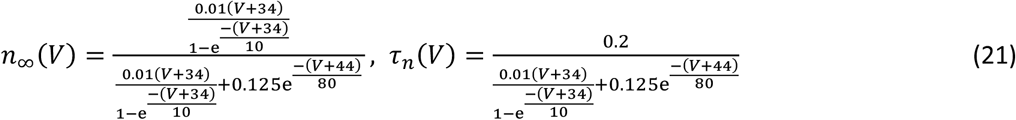

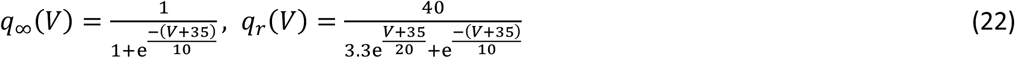

The INT received excitatory input from the PYR, with conductance *g_ie_*, reversal potential *E_e_*, and synaptic variable 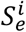; and inhibitory input from itself, with the synaptic conductance *g_ii_* reversal potential *E_i_* and synaptic variable 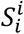. The synaptic variables were modeled using the same formalism as the inhibitory synapse on the PYR, namely

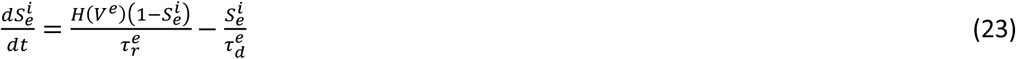

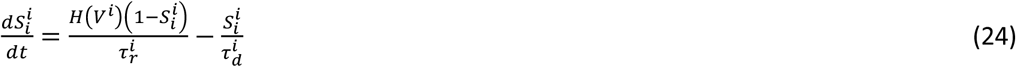

with the same sigmoid activation function as for the inhibitory synapse on the PYR (**equation 17**). For modeling the INT-to-PYR network without gamma resonance on the INT (**Fig. 7B**), 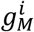 was set to zero. For modeling the INT with gamma resonance but without the PYR (**Fig. 7C**), *g_ei_* and *g_ii_* were set to zero. The full model was used for **Fig. 7D**.

## REFERENCES

Bi GQ, Poo MM (2001) Synaptic modification by correlated activity: Hebb’s postulate revisited. Annu. Rev. Neurosci. 24:139–66.

Boehlen A, Heinemann U, Henneberger C (2011) Heterogeneous voltage dependence of interneuron resonance in the hippocampal stratum radiatum of adult rats. Synapse, 65:1378–81.

Brown DA, Adams PR (1980) Muscarinic suppression of a novel voltage-sensitive K+ current in a vertebrate neurone. Nature 283:673–6.

Buhl EH, Halasy K, Somogyi P (1994) Diverse sources of hippocampal unitary inhibitory postsynaptic potentials and the number of synaptic release sites. Nature 368:823–8.

Chapman CA, Lacaille JC (1999) Intrinsic theta-frequency membrane potential oscillations in hippocampal CA1 interneurons of stratum lacunosum-moleculare. J. Neurophsyiol. 81:1296–307.

Chen Y, Li X, Rotstein HG, Nadim F (2016) Membrane potential resonance frequency directly influences network frequency through electrical coupling. J. Neurophysiol. 116:1554–63.

Cobb SR, Buhl EH, Halasy K, Paulsen O, Somogyi P (1995) Synchronization of neuronal activity in hippocampus by individual GABAergic interneurons. Nature 378:75–8.

Dayan P, Abbott LF (2001) Theoretical neuroscience. Cambridge, Massachusetts: MIT Press.

Drover JD, Tohidi V, Bose A, Nadim F (2007) Combining synaptic and cellular resonance in a feed-forward neuronal network. Neurocomputing 70:2041–5.

Engel TA, Schimansky-Geier L, Herz AV, Schreiber S, Erchova I (2008) Subthreshold membranepotential resonances shape spike-train patterns in the entorhinal cortex. J. Neurophysiol. 100:1576–88.

Fox DM, Rotstein HG, Nadim F (2016) Neuromodulation produces complex changes in resonance profiles of neurons in an oscillatory network. SFN Abstract 811.08.

Fries P (2015) Rhythms for cognition: communication through coherence. Neuron 88:220–35.

Fujisawa S, Amarasingham A, Harrison MT, Buzsáki G (2008) Behavior-dependent short-term assembly dynamics in the medial prefrontal cortex. Nat. Neurosci. 11:823–33.

Gutfreund Y, Yarom Y, Segev I (1995) Subthreshold oscillations and resonant frequency in guinea-pig cortical neurons: physiology and modelling. J. Physiol. 483:621–40.

Helfrich RF, Breska A, Knight RT (2019) Neural entrainment and network resonance in support of top-guided attention. Curr. Opin. Psychol. 29:82–9.

Hodgkin AL, Huxley AF (1952) A quantitative description of membrane current and its application to conductance and excitation in nerve. J. Physiol. 117:500–44.

Hu H, Vervaeke K, Storm JF (2002) Two forms of electrical resonance at theta frequencies, generated by M-current, h-current and persistent Na+ current in rat hippocampal pyramidal cells. J. Physiol. 545:783–805.

Hu H, Vervaeke K, Graham LJ, Storm JF (2009) Complementary theta resonance filtering by two spatially segregated mechanisms in CA1 hippocampal pyramidal neurons. J. Neurosci. 29:14472–83.

Hutcheon B, Miura RM, Puil E (1996) Subthreshold membrane resonance in neocortical neurons. J. Neurophysiol. 76:683–97.

Hutcheon B, Yarom Y (2000) Resonance, oscillation and the intrinsic frequency preferences of neurons. Trends Neurosci. 23:216–22.

Iaccarino HF, Singer AC, Martorell AJ, Rudenko A, Gao F, Gillingham TZ, Mathys H, Seo J, Kritskiy O, Abdurrob F, Adaikkan C, Canter RG, Rueda R, Brown EN, Boyden ES, Tsai LH (2016) Gamma frequency entrainment attenuates amyloid load and modifies microglia. Nature 540:230–35

Izhikevich EM, Desai NS, Walcott EC, Hoppensteadt FC (2003) Bursts as a unit of neural information: selective communication via resonance. Trends Neurosci. 26:161–7.

Izhikevich EM (2007) Dynamical systems in neuroscience. Cambridge, Massachusetts: MIT Press.

Kandel ER, Spencer WA (1961) Electrophysiology of hippocampal neurons. II. After-potentials and repetitive firing. J. Neurophysiol. 24:243–59.

Kang YJ, Lewis HES, Young MW, Govindaiah G, Greenfield Jr. LJ, Garcia-Rill E, Lee SH (2018) Cell type-specific intrinsic perithreshold oscillations in hippocampal GABAergic interneurons. Neuroscience, 376:80–93.

Lakatos P, Karmos G, Mehta AD, Ulbert, I, Schroeder CE (2008) Entrainment of neuronal oscillations as a mechanism of attentional selection. Science 320:110–3.

Lampl I, Yarom Y (1997) Subthreshold oscillations and resonant behaviour: Two manifestations of the same mechanism. Neuroscience 78:325–41.

Ledoux E, Brunel N (2011) Dynamics of networks of excitatory and inhibitory neurons in response to time-dependent inputs. Front. Comput. Neurosci. 5:25.

Leiser RJ, Rotstein HG (2019) Network resonance: impedance interactions via a frequency response alternating map (FRAM). SIAM J. Appl. Dyn. Syst. 18:769–807.

Lisman J (2005) The theta/gamma discrete phase code occuring during the hippocampal phase precession may be a more general brain coding scheme. Hippocampus 15:913–22.

Llinás R, Jahnsen H (1982) Electrophysiology of mammalian thalamic neurones in vitro. Nature 297:406–8.

Llinás R, Yarom Y (1986) Oscillatory properties of guinea-pig inferior olivary neurones and their pharmacological modulation: an in vitro study. J. Physiol. 376:163–82.

Markram H, Wang Y, Tsodyks M (1998) Differential signaling via the same axon of neocortical pyramidal neurons. Proc. Natl. Acad. Sci. USA 95:5323–8.

Pervouchine DD, Netoff TI, Rotstein HG, White JA, Cunningham MO, Whittington MA, Kopell NJ (2006) Low-dimensional maps encoding dynamics in entorhinal cortex and hippocampus. Neural Comp. 18:2617–50.

Pike FG, Goddard RS, Suckling JM, Ganter P, Kasthuri N, Paulsen O (2000) Distinct frequency preferences of different types of rat hippocampal neurones in response to oscillatory input currents. J. Physiol. 529:205–13.

Poolos NP, Migliore M, Johnston D (2002) Pharmacological upregulation of h-channels reduces the excitability of pyramidal neuron dendrites. Nat. Neurosci. 5:767–74.

Puil E, Gimbarzevsky B, Miura RM (1986) Quantification of membrane properties of trigeminal root ganglion neurons in guinea pigs. J. Neurophysiol. 55:995–1016.

Richardson MJ, Brunel N, Hakim V (2003) From subthreshold to firing-rate resonance. J. Neurophysiol. 89:2538–54.

Rotstein HG (2014) Frequency Preference Response to Oscillatory Inputs in Two-dimensional Neural Models: A Geometric Approach to Subthreshold Amplitude and Phase Resonance. J. Math. Neurosci. 4:11.

Rotstein HG, Nadim F (2014) Frequency preference in two-dimensional neural models: a linear analysis of the interaction between resonant and amplifying currents. J. Comput. Neurosci. 37:9–28.

Rotstein HG (2015) Subthreshold amplitude and phase resonance in models of quadratic type: nonlinear effects generated by the interplay of resonant and amplifying currents. J. Comput. Neurosci. 38:325–54.

Rotstein HG (2017a) The shaping of intrinsic membrane potential oscillations: positive/negative feedback, ionic resonance/amplification, nonlinearities and time scales. J. Comput. Neurosci. 42:133–66.

Rotstein HG (2017b) Resonance modulation, annihilation and generation of anti-resonance and anti-phasonance in 3D neuronal systems: interplay of resonant and amplifying currents with slow dynamics. J. Comput. Neurosci. 43:35–63.

Rotstein HG (2017c) Spiking resonances in models with the same slow resonant and fast amplifying currents but different subthreshold dynamic properties. J. Comput. Neurosci. 43:243–71.

Rotstein HG, Ito T, Stark E (2017) Inhibition-based theta spiking resonance in a hippocampal network. SFN Abstract 615.11.

Sirota A, Montgomery S, Fujisawa S, Isomura Y, Zugaro M, Buzsáki G (2008) Entrainment of neocortical neurons and gamma oscillations by the hippocampal theta rhythm. Neuron 80:683–97.

Smeal RM, Ermentrout GB, White JA (2010) Phase-response curves and synchronized neural networks. Philos. Trans. R. Soc. Lond. B Biol. Sci. 365:2407–22.

Stark E, Eichler R, Roux L, Fujisawa S, Rotstein HG, Buzsáki G (2013) Inhibition-induced theta resonance in cortical circuits. Neuron 80:1263–76.

Stark E, Roux L, Eichler R, Senzai Y, Royer S, Buzsáki G (2014) Pyramidal cell-interneuron interactions underlie hippocampal ripple oscillations. Neuron 83:467–80.

Stark E, Roux L, Eichler R, Buzsáki G (2015) Local generation of multi-neuronal spike sequences in the hippocampal CA1 region. Proc. Natl. Acad. Sci. USA, 112:10521–6.

Thomson AM, Deuchars J, West DC (1993) Large, deep layer pyramid-pyramid single axon EPSPs in slices of rat motor cortex display paired pulse and frequency-dependent depression, mediated presynaptically and self-facilitation, mediated postsynaptically. J. Neurophysiol. 70:2354–69.

Vierling-Claassen D, Siekmeier P, Stufflebeam S, Kopell N (2008) Modeling GABA alterations in schizophrenia: a link between impaired inhibition and altered gamma and beta range auditory entrainment. J. Neurophysiol. 99:2656–71.

Wang XJ, Buzsáki G (1996) Gamma oscillation by synaptic inhibition in a hippocampal interneuronal network model. J. Neurosci. 16:6402–13.

Wang XJ (2010) Neurophysiological and computational principles of cortical rhythms in cognition. Physiol. Rev. 90:1195–268.

Whittington MA, Traub RD, Jefferys JG (1995) Synchronized oscillations in interneuron networks driven by metabotropic glutamate receptor activation. Nature 373:612–5.

Zhou Y, Vo T, Rotstein HG, McCarthy MM, Kopell N (2018) M-current expands the range of gamma frequency inputs to which the neuronal target entrains. J. Math. Neurosci. 8:13.

Zucker RS (1989) Short-term synaptic plasticity. Annu. Rev. Neurosci. 12:13–31.

Zucker RS, Regehr WG (2002) Short-term synaptic plasticity. Annu. Rev. Physiol. 64:355–405.

## REFERENCES FOR METHODS

Borgers C, Talei Franzesi G, Lebeau FE, Boyden ES, Kopell NJ (2012) Minimal size of cell assemblies coordinated by gamma oscillations. PLoS Comput. Biol. 8:e1002362.

Dayan P, Abbott LF (2001) Theoretical neuroscience. Cambridge, Massachusetts: MIT Press.

Ermentrout GB, Kopell N (1998) Fine structure of neural spiking and synchronization in the presence of conduction delays. Proc. Natl. Acad. Sci. USA 95:1259–64.

Ermentrout GB, Terman D (2010) Mathematical Foundations of Neuroscience. Springer, Berlin.

Fee MS, Mitra PP, Kleinfeld D (1996) Automatic sorting of multiple unit neuronal signals in the presence of anisotropic and non-Gaussian variability. J. Neurosci. Methods 69:175–88.

Gaspar N, Eichler R, Stark E (2019) A novel low-noise movement tracking system with real-time analog output for closed-loop experiments. J. Neurosci. Methods, 318:69–77.

Harris KD, Henze DA, Csicsvari J, Hirase H, Buzsáki G (2000) Accuracy of tetrode spike separation as determined by simultaneous intracellular and extracellular measurements. J. Neurophysiol. 84:401–14.

Manor Y, Nadim F (2001) Synaptic depression mediates bistability in neuronal networks with recurrent inhibitory connectivity. J. Neurosci. 21:9460–70.

Olufsen MS, Whittington MA, Camperi M, Kopell N (2003) New roles for the gamma rhythm: population tuning and preprocessing for the Beta rhythm. J. Comput. Neurosci. 14:33–54.

Pachitariu M, Steinmetz NA, Kadir SN, Carandini M, Harris KD (2016) Fast and accurate spike sorting of high-channel count probes with KiloSort. In: Advances in Neural Information Processing Systems 29 (Ed: Lee DD, Sugiyama M, Luxburg UV, Guyon I, Garnett R), 4448–56.

Schmitzer-Torbert N, Jackson J, Henze D, Harris K, Redish AD (1996) Quantitative measures of cluster quality for use in extracellular recordings. Neuroscience 131:1–11.

Stark E, Abeles M (2009) Unbiased estimation of precise temporal correlations between spike trains. J. Neurosci. Methods 179:90–100.

Stark E, Koos T, Buzsáki G (2012) Diode probes for spatiotemporal optical control of multiple neurons in freely moving animals. J. Neurophysiol. 108:349–63.

Zemankovics R, Káli S, Paulsen O, Freund TF, Hájos N (2010) Differences in subthreshold resonance of hippocampal pyramidal cells and interneurons: the role of h-current and passive membrane characteristics. J. Physiol. 588:2109–32.

